# SCF^cyclin F^ - EXO1 axis controls cell cycle dependent execution of double strand break repair

**DOI:** 10.1101/2023.10.26.562393

**Authors:** Hongbin Yang, Shahd Fouad, Paul Smith, Eun Young Bae, Yu Ji, Ava Van Ess, Francesca M Buffa, Roman Fisher, Iolanda Vendrell, Benedikt M Kessler, Vincenzo D’Angiolella

## Abstract

Ubiquitination is a crucial post-translational modification required for the proper repair of DNA double-strand breaks (DSBs) induced by ionising radiation (IR). DSBs are mainly repaired through homologous recombination (HR) when template DNA is present and non-homologous end joining (NHEJ) in its absence. Additionally, microhomology-mediated end joining (MMEJ) and single strand annealing (SSA) provide back-up DSBs repair pathways. However, the mechanisms controlling their use remain poorly understood. By employing a high-resolution CRISPR screen of the ubiquitin system after IR, we systematically uncover genes required for cell survival and elucidate a critical role of the E3 ubiquitin ligase SCF^cyclin^ ^F^ in cell cycle-dependent DSB repair. We show that SCF^cyclin^ ^F^ -mediated EXO1 degradation prevents DNA end resection in mitosis, allowing MMEJ to take place. Moreover, we identify a conserved cyclin F recognition motif, distinct from the one used by other cyclins, with broad implications in cyclin specificity for cell cycle control.

## Introduction

The maintenance of genomic integrity is a fundamental aspect of cellular homeostasis for cell survival. Among the different DNA lesions, double-strand breaks (DSBs) pose a significant threat to genome stability, as they or their improper repair can lead to loss of genetic information, chromosomal rearrangements, oncogenic transformations, and cell death. To safeguard against these deleterious outcomes, cells have evolved an intricate and tightly regulated DNA damage response (DDR) network, including multiple DNA damage repair pathways. Homologous recombination (HR) and non-homologous end joining (NHEJ) are the major DSB repair pathways in mammalian cells. NHEJ joins the DNA ends with minimal sequence homology in an error-prone manner, while HR utilizes the sister chromatid only available in late S/G2 phases of the cell cycle as a template for error-free repair.

A key regulatory process in the DDR network is ubiquitination, a covalent and reversible post-translational modification mediated by the E1-E2-E3 cascade (ubiquitin-activating enzyme, ubiquitin-conjugating enzyme, and ubiquitin ligase, respectively). Ubiquitination orchestrates the recruitment of DNA repair factors, signal transduction, and the temporal control of DNA repair pathways ^1^. Moreover, ubiquitination is also known to regulate the pathway choice between HR and NHEJ^2, 3^. Besides these major repair pathways, microhomology-mediated end joining (MMEJ) and single-strand annealing (SSA) are considered back-up mechanisms when HR and NHEJ are compromised ^4^. In contrast to the extensively studied HR and NHEJ pathways, our knowledge of how MMEJ and SSA are executed and regulated remains scarce; they are simply known to be error-prone and characteristically dependent on DNA polymerase theta (POLQ) and RAD52, respectively ^5^. HR and SSA both require exonuclease 1 (EXO1)- or DNA2-mediated long-range 5’→3’ nucleolytic degradation and the sequential generation of a 3’ single-stranded DNA overhang— a process known as the long-range DNA end resection ^6^. As EXO1- or DNA2-mediated DNA end resection can remove several hundreds to a few thousands nucleotides^7, 8^, the activities of these exonucleases need to be tightly regulated and precisely executed to avoid DNA breakage during damage repair, loss of genetic information, and, to an even worse extent, unwanted recombination due to extensive homology exposure among random genomic regions. An example of such regulation is the post-translational modification or direct inhibition of EXO1 upon DNA damage induction ^9–12^. DNA structures produced by long-range end resection are known to be poor substrates for NHEJ and MMEJ machineries, indicating a possible role of DNA end resection in dictating DSB repair pathway choice. To date, various mechanisms have been reported to direct pathway choice between NHEJ and HR^13, 14^. However, the role of ubiquitination in DNA end resection and in determining alternative DSB repair pathways choice is understudied.

Cyclin F is the funding member of the F-box family proteins, the characteristic F-box domains of which can interact with the SKP1-CUL1 complex so that the F-box protein can function as a substrate receptor (the direct substrate interactor) in the SKP1-CUL1-F-box (SCF) E3 ligases holocomplex. Approximately 70 F-box proteins have been identified in mammal. SCF complex that utilises cyclin F as the substrate receptor is called the SCF^cyclin^ ^F^ E3 ligase. SCF^cyclin^ ^F^ was previously reported to degrade various proteins involved in cell cycle progression and in DNA damage repair ^15–18^. Starting from a focused CRISPR screen to systematically discover E1s, E2s, and E3s that are required for cell survival after ionising radiation (IR), we identified *CCNF* (the gene encoding cyclin F) as a crucial determinant of cell survival after IR. The following mechanistic study revealed that SCF^cyclin^ ^F^ degrades EXO1 in the G2 and M cell cycle phases to prevent DNA hyper-resection and SSA, and promote MMEJ. Disruption of this axis either by *CCNF* knockout or by expression of a non-degradable R842A EXO1 mutant promotes hyper-resection and chromosome aberrations, which in turn increases IR-induced cell death. This study unravels a novel mechanism to regulate DSB back-up repair pathways in the context of cell cycle.

## Results

### CRISPR screen of the ubiquitin system identifies genes required for cell survival after IR

Radiotherapy by ionizing radiation (IR) is used extensively to treat cancer in both palliative and curative contexts. For some tumours, such as glioblastoma multiforme (GBM), radiotherapy remains the major therapeutic strategy besides surgery due to the scarcity of chemotherapy options. However, patients still suffer from dismal prognoses due to GBM cells’ fast development of radioresistance. It is, therefore, imperative to identify the major pathways conferring IR resistance so as to develop strategies to re-sensitize cancer cells to radiation. To this end, we performed an unbiased CRISPR screen on genes involved in ubiquitin conjugation for the identification of radioprotective genes in GBM cells. We tested the IR sensitivity of 7 glioblastoma-derived cell lines via colony formation assay following IR (Figure S1A) and identified LN229 as the most radioresistant, and therefore the best model for drop-out screens— a screen to identify genes whose depletion promotes radiosensitivity.

Cas9 was introduced into the LN229 cells using a lentiviral expression system and a clonal population was then generated to achieve a high level of Cas9 expression (Figure S1B). To minimize variations caused by Cas9 expression and clonal effects, we isolated several clones and selected the one that showed no differences in cell proliferation and radio-sensitivity when compared to the LN229 parental cells (Figure S1C).

A commercially available library focused on the ubiquitin system was utilized in the screen. The library covers 5 E1s, 35 E2s, 531 established or putative E3s, 31 core essential genes as positive controls, and 100 non-targeting controls. Each gene was targeted with 10 sgRNA guides and we chose a fold representation of ∼750 in order to gain more resolution in the screen (Figure 1A). Cells were treated with 4 Gy IR in a single dose, which was previously found to kill around 50% of LN229 cells via colony formation assays (Figure S1A and S1C).

**Figure 1.**
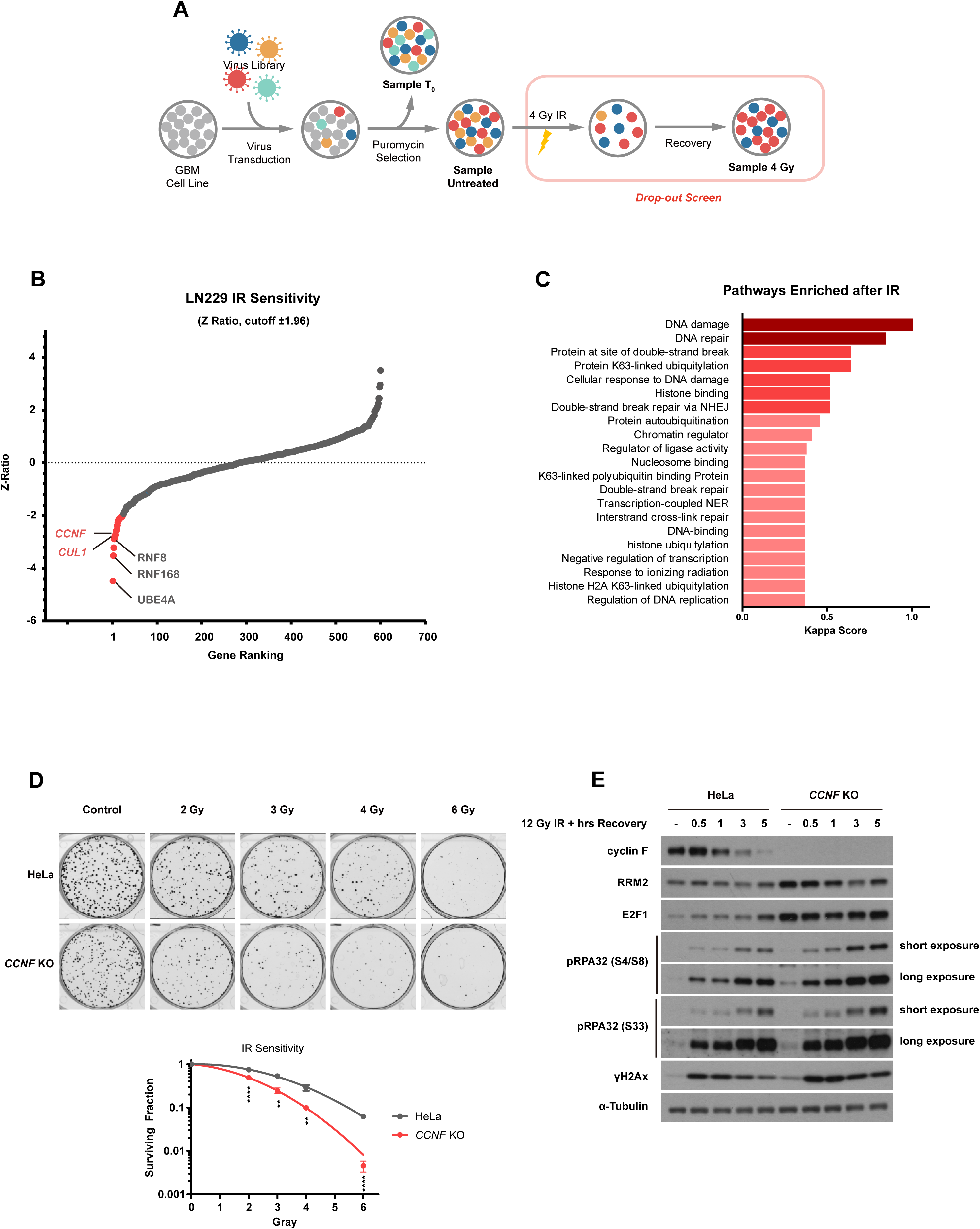
CRISPR screen of the ubiquitin system identified genes required for cell survival after IR. A. Schematic representation of the CRISPR screen. Two days after transducing Cas9-expressing LN229 with the sgRNA library, Sample T_0_ was collected as a reference sample of sgRNA expression. Cells were then selected with puromycin for 7 days before ionising radiation (IR) treatment. 14 days after IR, cells were collected for genomic DNA extraction and PCR amplification of the sgRNA sequences. PCR product was sent for next generation sequencing (NGS) so that change in relative abundance of each sgRNA before and after IR can be assessed. B. Identification of genes impacting IR sensitivity in LN229 by Z-Ratio. C. Over-representation analysis (ORA) of statistically significant hits identified from the screen (absolute value of Z-Ratio>1.96). Pathways coloured in dark red are highly confident (κ>1). Pathways in red are confident (κ>0.5). Pathways in pink are potential pathways (κ<0.5). D. Colony formation assay in *CCNF* K/O’s cells. HeLa parental cells or HeLa *CCNF* K/O cells were seeded for colony formation assay and challenged with the indicated dose of IR. 7 days after IR, cells were stained with crystal violet and counted. Error bars represent standard deviations of three biological replicates. Statistical analysis two-tailed unpaired t test. ** indicates P ⩽ 0.01, *** indicates P ⩽ 0.001, and **** indicates P ⩽ 0.0001. E. Immunoblotting to detect responses to IR treatment as indicated by DNA damage markers— pRPA32 S4/8 (single stranded DNA at double strand break), pRPA32 S33 (single strand breaks), and γH2Ax (DNA double strand breaks amount).

The replicates of the screen showed significant correlation and clustering in the principal component analysis (PCA) (Figure S1D and S1E). In addition, we calculated the effect size for all guides using Cohen D ^19^(Figure S1F). The score indicates a large effect size, highlighting the high sensitivity and resolution of the screen.

We first analysed the results in untreated cells, comparing cells harvested immediately after library transduction (Sample T_0_) to cells that survived puromycin selection (Sample Untreated). We not only identified positive control genes (core essential genes) provided in the library but also a set of essential genes specific to LN229 (Figure S1G). Altogether, these results indicate a good retrieval of essential genes.

To identify the genes required for survival after IR, we compared cells treated with 4 Gy IR to the untreated cells. A Z-ratio score was plotted using CRISPRAnalyzeR (http://crispr-analyzer.dkfz.de) (Figure 1B). Over-representation analysis (ORA) was performed on the statistically significant genes that conferred either IR sensitivity or IR resistance (with Z-ratio > 1.96 or < −1.96). A significant enrichment was detected for genes participating in DNA damage and repair (Figure 1C).

It is worth noting that we identified *UBE4A*, *RNF168* and *RNF8* as top hits. These are all genes known to have crucial roles in mediating DNA damage responses and repair ^20–22^, indicating the screen could also potentially reveal hitherto uncharacterized players in these processes. We also identified *UBA3*, the E1 for neddylation Neddylation inhibitor MLN4924 is a potent radiosensitiser with an unknown mechanism, and our finding further supports the idea that neddylation is required in responding to and repairing DNA ^23–25^. In addition to these well-established DNA damage regulators, we also identified CUL1, the scaffolding protein of SCF complexes, and several F-box proteins, which function as substrate receptors in SCFs.

Among the F-box adaptors identified from the screen, *CCNF* ranks the highest, making it a good target for validation. To this end, we employed a classic colony formation assay in HeLa cells where the *CCNF* gene has been knocked out using CRISPR. *CCNF* knockout (*CCNF* K/O) significantly sensitised HeLa cells to IR at all doses tested (Figure 1D).

We also performed straight western blotting (WB) to assess the DDR. *CCNF* K/O significantly upregulated the steady-state levels of the previously reported SCF^cyclin^ ^F^ substrates —RRM2 ^15^ and E2F1 ^17^. In addition, the delivery of IR to *CCNF* K/O cells also induced upregulation of RPA32 phosphorylation at Ser4/Ser8 (a marker of single-strand DNA (ssDNA) at DSB repair sites upon activation of signalling proteins ATM and DNA-PK) and Ser33 (marker of ssDNA upon ATR activation) at steady-state levels and after IR. Moreover, the classic DSB marker γH2Ax was also upregulated in *CCNF* K/O cells compared to the parental line, both at steady state and after IR (Figure 1E), indicating that loss of cyclin F could increase DNA damage at a basal level and prevent or delay the repair of damaged DNA after IR treatment.

### Cyclin F interacts with EXO1 during G2 and M and after IR, facilitating its ubiquitination

To understand how cyclin F would regulate IR sensitivity and DNA damage response and repair, we established a cell line expressing cyclin F fused to TurboID (an engineered biotin ligase that conjugates biotin to proteins within tens of nanometres of its fused protein) to identify the cyclin F interactome (Figure S2A). This approach, which exploits a non-specific biotin ligase, has been used extensively to identify interacting partners of cellular proteins ^26^, with improvements made to enhance specificity and efficiency ^27^. We compared untreated cells to those treated with MLN4924 to generate a comprehensive view of all potential SCF^cyclin^ ^F^ substrates (Figure 2A). MLN4924 helps to enrich cyclin F-substrate interactions by inhibiting the ligation of ubiquitin to substrates, which would then cause dissociation of the substrate from cyclin F. As shown in Figure 2A, several proteins are significantly enriched following MLN4924 treatment, with EXO1 identified as one of the top hits.

**Figure 2.**
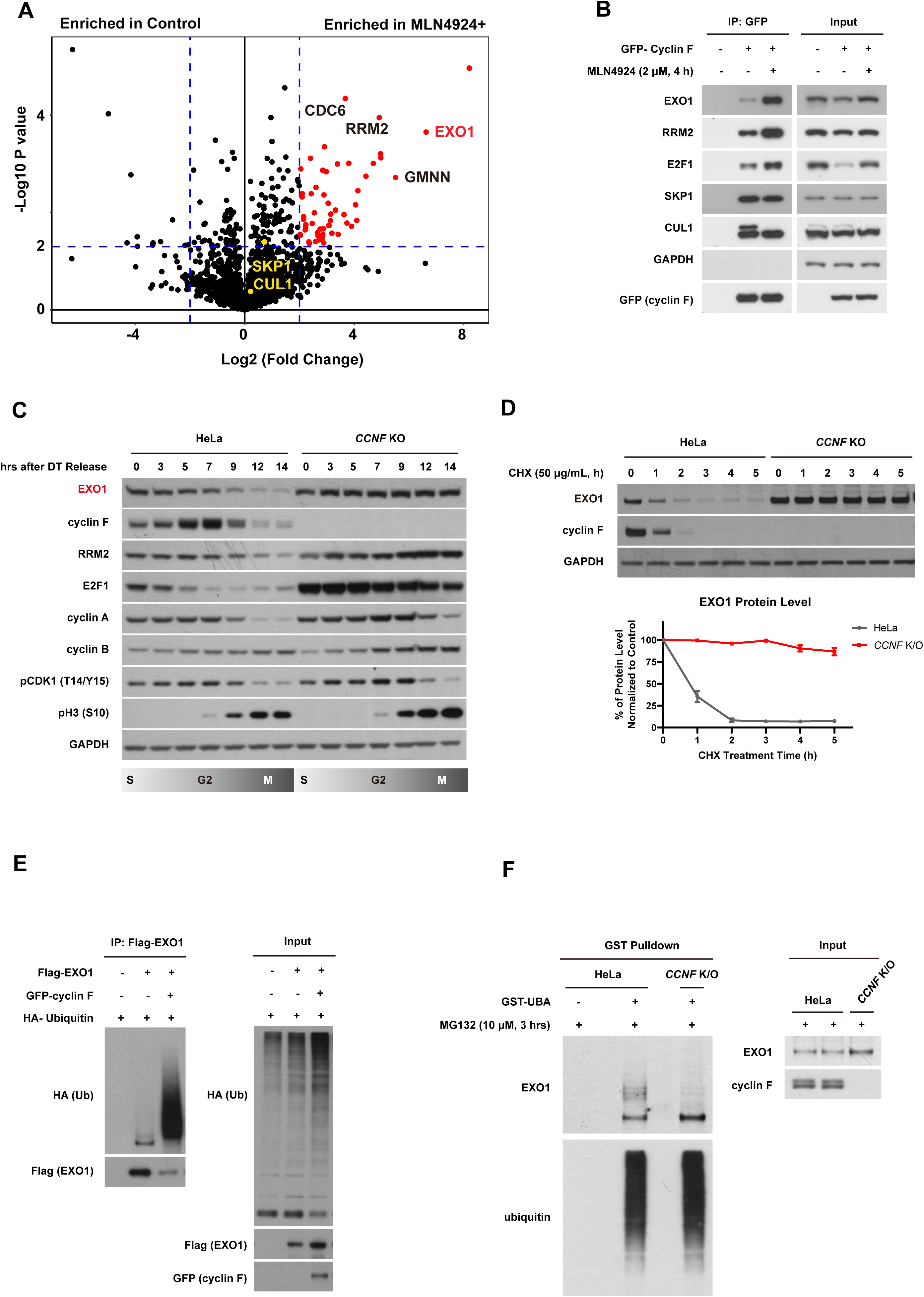
Cyclin F interacts and SCF^cyclin^ ^F^ ubiquitinates EXO1. A. Volcano plot representing mass spectrometry analysis of differential TurboID-cyclin F interacting partners in the presence *vs* absence of MLN4924. Proteins were isolated with streptavidin after labelling for 1 hour with biotin. B. GFP-tagged cyclin F was expressed in HEK293T before immunoprecipitation *via* GFP agarose beads. Immunoprecipitated material was analysed by immunoblotting as indicated. C. Immunoblotting of cell cycle synchronised HeLa cells *via* double thymidine (DT) block-release. D. Immunoblotting after treating cells with Cycloheximide (CHX) for the indicated time -*upper panel*. Relative quantification of EXO1 protein levels in HeLa parental cells or *CCNF* K/O cells after normalisation with EXO1 levels at T0 for each cell line - *bottom panel*. Error bars: standard deviations of three biological replicates; E. Immunoblotting after expression of GFP-cyclin F, Flag-EXO1, and HA-ubiquitin in HEK293T. Flag-EXO1 is isolated *via* Flag agarose beads pulldown before immunoblotting. F. Immunoblotting after isolation of endogenous ubiquitinated proteins using recombinant GST-tagged UBA domain of UBIQLN protein. Input samples before immunoprecipitation are indicated – *right panel*.

Interaction between cyclin F and EXO1 was then validated via immunoprecipitation-western blotting. As shown in Figure 2B, with complete abrogation of CUL1 neddylation as validation of MLN4924 treatment’s efficacy, MLN4924 promoted cyclin F interaction not only with the well-established substrates RRM2 and E2F1, but also with EXO1. On the contrary, interaction between cyclin F and the SCF components CUL1 and SKP1 remained intact (Figure 2B).

As cyclin F oscillates during the cell cycle and only peaks in G2, cyclin F substrates are expected to be degraded mainly in the G2 and M phases^15–17^. To test this, cell cycle synchronisation via double thymidine block release was performed in HeLa parental and *CCNF* K/O cells to assess changes in EXO1 protein levels from the start of the G1/S transition to mitosis. As shown in Figure 2C, EXO1 protein levels started to decrease 7 hours after cells were released into thymidine-free media — the exact time when cyclin F level peaks in G2. EXO1 is almost entirely depleted at the 14-hour time point when cells enter mitosis (evidenced by two markers of CDK1 activation - phosphorylation of histone H3 on serine 10 and loss of phosphorylated CDK1 on threonine 14 / tyrosine 15). However, this downregulation of EXO1 in G2 and M can be completely abrogated by *CCNF* K/O (Figure 2C).

To further test whether EXO1 is a SCF^cyclin^ ^F^ substrate targeted for degradation, we performed a cycloheximide (CHX) chase experiment. As CHX is a ribosomal toxin and can block protein synthesis, net degradation and therefore half-lives of proteins can be assessed in CHX chase experiments. As shown in Figure 2D, the half-life of EXO1 is less than 1 hour in parental cells but increases to over 5 hours in *CCNF* K/O cells (Figure 2D). Similar results were also obtained with si*CCNF* in LN229 cells (Figure S2B). Conversely, over-expressed cyclin F was found to downregulate EXO1 protein level in both LN229 and HEK293T. This down-regulation is rescued by the proteasome inhibitor MG132 (Figure S2C), suggesting cyclin F-mediated EXO1 downregulation is proteasome-dependent and further supporting the idea that EXO1 is a SCF^cyclin^ ^F^ substrate targeted for proteasomal degradation.

To establish a direct role of cyclin F in ubiquitinating and degrading EXO1, we employed two methods to detect EXO1 ubiquitination. We first isolated overexpressed EXO1 proteins following cyclin F overexpression in HEK293T and observed a clear upregulation of EXO1 ubiquitination (Figure 2E). Overexpressed ubiquitin and E3 ligases can lead to artefacts, however, so we then probed EXO1 ubiquitination at an endogenous level. After unbiasedly enriching all ubiquitinated proteins from parental HeLa cells using the recombinant ubiquitin-associated domain (UBA domain) of the UBIQLN protein in an ubiquitin binding entity (UBE) pulldown assay ^28^, we managed to detect endogenous levels of polyubiquitinated EXO1 in HeLa parental cells using an EXO1 antibody (Figure 2F). More importantly, this polyubiquitinated smear signal is completely lost in the *CCNF* K/O cells, suggesting SCF^cyclin^ ^F^ as a major E3 responsible for the ubiquitination of endogenous EXO1 (Figure 2F).

To further elucidate the chain specificity of SCF^cyclin^ ^F^-mediated EXO1 ubiquitination, we performed a ubiquitination assay by overexpressing either wild-type (WT) ubiquitin or K48R ubiquitin mutant. As shown in Figure S2D, the K48R mutation completely abrogated both basal-level EXO1 ubiquitination and cyclin F overexpression-induced EXO1 ubiquitination, demonstrating that EXO1 ubiquitination uses K48 linkages.

We observed reduced levels of EXO1 3 hours after IR treatment in accordance with findings by Tomimatsu et al ^9^. This reduction of EXO1 is also fully cyclin F-dependent because in *CCNF* K/O cells, we could detect high levels of EXO1 after IR (Figure S3A). We then measured the interaction between EXO1 and cyclin F at different time points after IR treatment and detected an increase over time, suggesting that cyclin F also promotes EXO1 degradation after IR (Figure S3B). We cannot exclude that the degradation of EXO1 in these conditions is triggered by cells moving from S phase to mitosis after IR, but the interaction between cyclin F and RRM2 was reduced in the same experiments, suggesting the presence of additional modes of EXO1 regulation after DNA damage. The cyclin F-EXO1 interaction was also promoted by treatment with camptothecin (CPT), a topoisomerase I inhibitor known to arrest cells in S phase (Figure S3C). Furthermore, endogenous ubiquitination of EXO1 was increased after treating cells with IR as detected by UBE assay. The ubiquitination ladder was fully dependent on cyclin F as it disappeared upon cyclin F depletion (Figure S3D).

Taken together, our findings demonstrate that SCF^cyclin^ ^F^ is the major E3 targeting EXO1 for K48-linked polyubiquitination both in mitosis and after DNA damage.

### EXO1 accumulation Leads to increased IR sensitivity upon cyclin F depletion

We have established that EXO1 is targeted for ubiquitination and degradation by SCF^cyclin^ ^F^ in the G2 and M phases of the cell cycle. However, the functional significance of this regulation remains unclear. Since EXO1 is one of the key exonucleases in DNA end resection after DSBs and participates in DNA damage repair, we speculated it could be the main substrate whose accumulation leads to IR hypersensitivity in *CCNF* K/O cells.

To test our hypothesis, we generated *EXO1* knockout (*EXO1* K/O) cells within a *CCNF* K/O background. In total, we generated four cell lines: HeLa WT, HeLa *EXO1* K/O, HeLa *CCNF* K/O, and HeLa *CCNF* K/O *EXO1* K/O (the double K/O), and assessed their sensitivity to IR. *CCNF* K/O showed increased sensitivity as shown previously (Figure 1D), while *EXO1* K/O cells did not have a significant impact on IR sensitivity (Figure 3A and 3B). In accordance with our hypothesis, the depletion of *EXO1* in a *CCNF* K/O background fully rescued the IR sensitivity observed in *CCNF* single knockouts, supporting the idea that EXO1 accumulation upon cyclin F loss is responsible for the IR hypersensitivity observed in *CCNF* K/O cells (Figure 3A and 3B). Moreover, DNA damage markers after IR endorsed this observation at a molecular level (Figure 3C), where elevated levels of RPA32 phosphorylation at S33 and S4/8 caused by *CCNF* K/O were fully rescued by *EXO1* K/O in the double K/O cell line. Similar results were obtained in LN229 using sgRNAs targeting *CCNF* and *EXO1* (Figure 3D). Taken together, these data demonstrate that the IR hypersensitivity observed upon cyclin F depletion is due to EXO1 accumulation.

**Figure 3.**
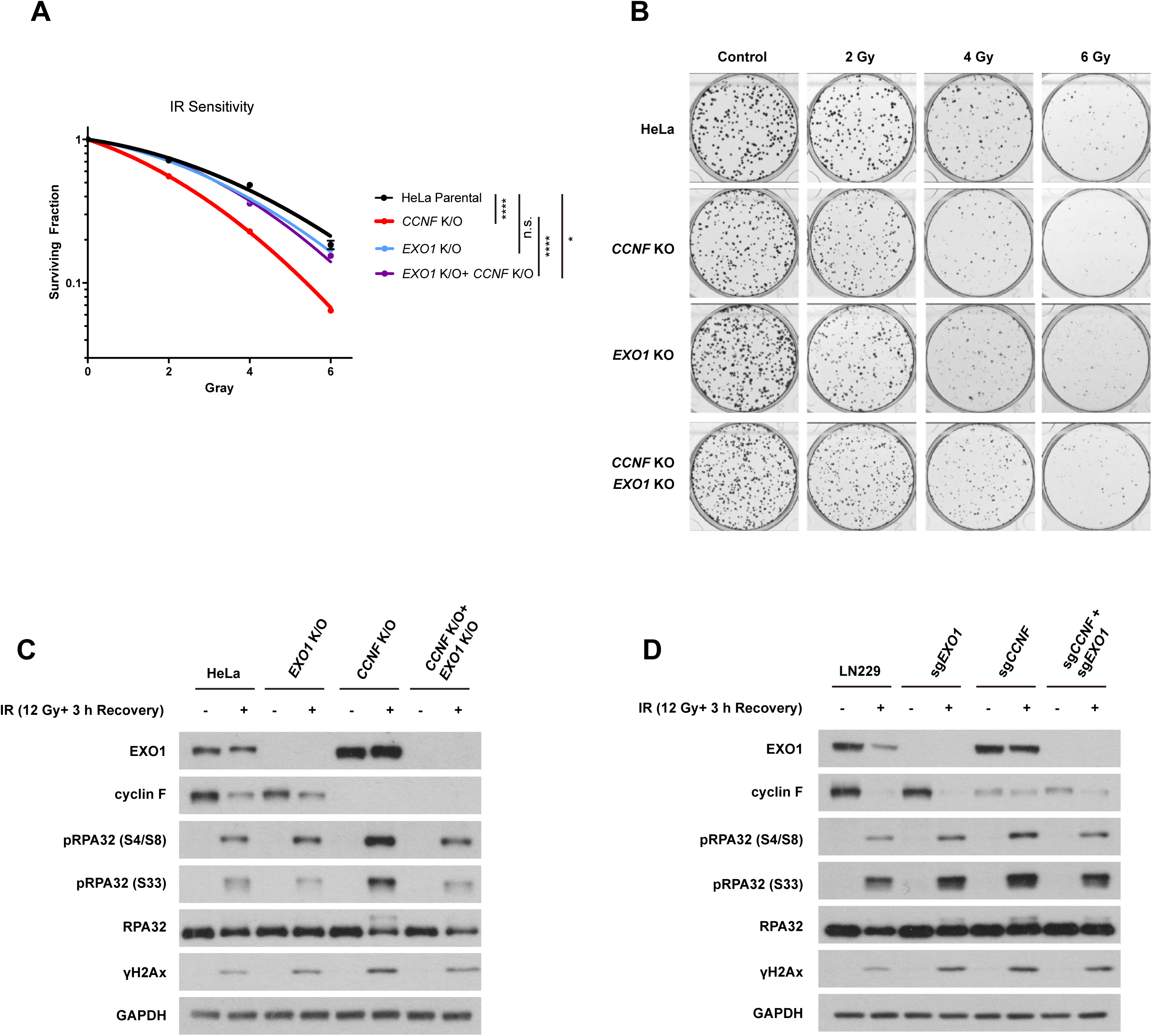
EXO1 accumulation leads to increased sensitivity to IR upon cyclin F depletion. A. HeLa parental cells, HeLa *CCNF* K/O, HeLa *EXO1* K/O and HeLa *CCNF* K/O *EXO1* K/O as indicated were seeded for colony formation assay and challenged with the indicated dose of IR. 7 days after IR, cells were stained with crystal violet and counted. Error bars represent standard deviations of three biological replicates. Statistical analysis two-tailed unpaired t test. ** indicates P ⩽ 0.01, *** indicates P ⩽ 0.001, and **** indicates P ⩽ 0.0001. B. Colony formation assay representative image of A. C. Immunoblotting to detect responses to IR treatment in HeLa cells as indicated by DNA damage markers. D. Immunoblotting to detect responses to IR treatment in LN229 as indicated by DNA damage markers. LN229 cells were transiently transfected with Cas9 protein and sgRNA as indicated 4 days before IR treatment.

### The RxIF (F-deg) motif defines a specific cyclin F recognition site

The mechanism of EXO1 recognition by cyclin F is not known, so we mapped the domains mediating the interactions on both proteins. We and others have previously shown that cyclin F uses the cyclin domain to recruit substrates ^29^. To establish that the cyclin domain is required for EXO1 recruitment and ubiquitination, we tested EXO1 interaction with two previously reported cyclin F mutants ^15, 16^ — ΔF (with L45A and P46A mutation in the F-box domain of cyclin F, compromising SKP1 and therefore CUL1 interaction) and ΔC (with M309A mutation, which disrupts the cyclin domain). Both wild-type cyclin F (WT) and ΔF were able to pull down endogenous EXO1, while the ΔC mutant was not (Figure 4A). *In vivo* ubiquitination experiments conducted with the same cyclin F mutants demonstrated a remarkable reduction in EXO1 ubiquitination following ΔF expression and a complete loss of ubiquitination if ΔC was expressed (Figure S4A). Taken together, these results demonstrate that cyclin F interacts with EXO1 through its cyclin domain.

**Figure 4.**
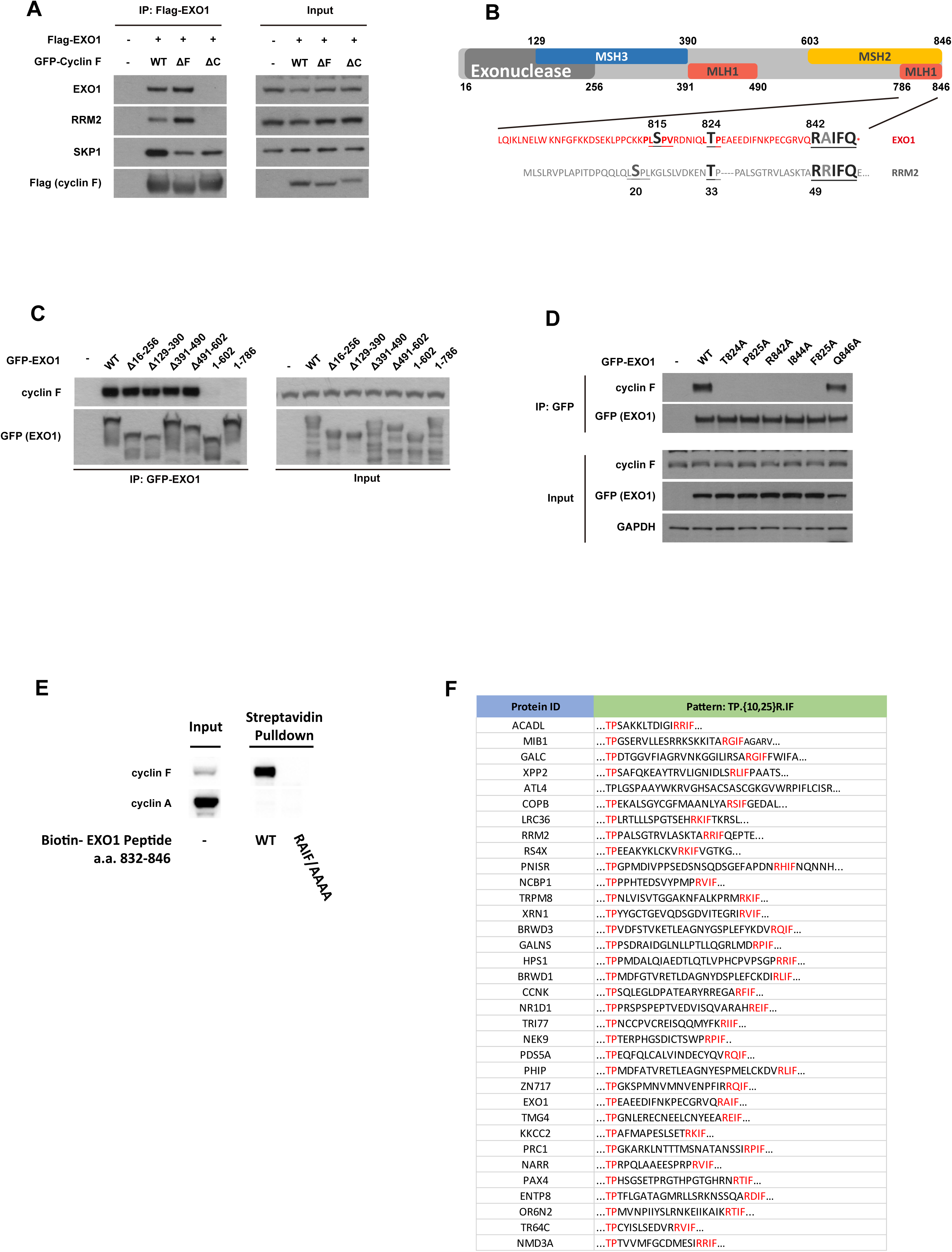
Cyclin Domain on Cyclin F and RAIF Motif on EXO1 are Required for Cyclin F-EXO1 Interaction. A. Immunoblotting after expression of GFP-cyclin F Wild Type (WT), GFP-cyclin F L35A/P36A (△F, which disrupts the F box domain), GFP-cyclin F M309A (△C, which disrupts the cyclin domain) with Flag-EXO1, and immunoprecipitation (IP) of Flag-EXO1 isolated *via* Flag agarose beads. B. Schematic representation of EXO1 depicting the localisation of the exonuclease domain, the MSH3 interaction domain, the MLH1 domains, and the MSH2 interaction domain. The last 60 amino acids of EXO1 C-terminus are highlighted and aligned with the previously characterised cyclin F interaction motif on RRM2. Amino acids presents in both EXO1 and RRM2 are enlarged and labelled in grey. C. Immunoblotting after immunoprecipitation (IP) of EXO1 WT and the indicated EXO1 fragments lacking the internal amino acids (Δ, x to y) or ending at 602 and 786. D. Immunoblotting after immunoprecipitation (IP) of GFP-EXO1 WT, GFP-EXO1 T824A, GFP-EXO1 P825A, GFP-EXO1 R842A, GFP-EXO1 I844A, GFP-EXO1 F845A and GFP-EXO1 Q846A mutants. E. Immunoblotting of pull down from cells using biotin-tagged EXO1 peptides encompassing the last 15 amino acid residues or the last 15 amino acid residues where RxIF was mutated to AAAA. F. Genome-wide identification of proteins containing an TP. R.IF. motif using SCANSITE 4.0.

Similarly, we mapped cyclin F interaction site(s) on EXO1. To this end, the exonuclease domain (amino acid residues from 16 to 256), MSH3 interaction domain (129-390), MLH1 interaction domain part 1 (391-490), MSH2 interaction domain (603-end), and MLH1 interaction domain part 2 (786-end) were isolated and tested for their interactions with endogenous cyclin F (domains are depicted in Figure 4B). As shown in Figure 4C, only C-terminal truncation on EXO1 disrupted its interaction with cyclin F, indicating the requirement of the last 60 amino acid residues on EXO1 for cyclin F interaction. Interestingly, alignment of the last 60 residues in EXO1 with the previously identified RRM2 degron revealed striking similarity in the two regions— a T824 residue in EXO1 closely resembles T33 in RRM2, which was previously identified as the priming phosphorylation site in initiating cyclin F binding, and an RxIFQ motif from 842 to 846 resembles residue 49 to 53 in RRM2, including an RRI motif previously identified as essential for cyclin F interaction ^15^. Since our findings potentially extended the well-established RxI motif to RxIFQ, we decided to test the essentiality of each of the RxIFQ residues in both RRM2 and EXO1. In both cyclin F substrates, the mutation of R, I and F to alanine prevented substrate interaction with cyclin F, while substitution of Q to A did not impact on cyclin F’s interaction with both EXO1 and RRM2 (Figure 4D and Figure S4B). As expected, an *in vivo* ubiquitination assay also demonstrated that mutations in the RxI motif abrogated ubiquitination (Figure S4C). To establish that RxIF was the essential motif for the interaction with cyclin F, we synthesized peptides containing the last 15 amino acids of EXO1 with the RxIF motif. This peptide was sufficient to retrieve cyclin F from cell lysates while mutations in the RxIF completely abolished its interaction with cyclin F (Figure 4E). Identification of this motif facilitated a homology-based search within the human proteome for other potential cyclin F substrates. The search retrieved 997 matches, including several well-established cyclin F substrates and interactors identified by LC/MS by us and other groups ^30–32^. Since the threonine site was also conserved in the cases of cyclin F interaction to both EXO1 and RRM2 and we confirmed the threonine’s essentiality in mediating cyclin F interaction to both substrates (Figure 4D), we restricted the search to proteins containing an RxIF motif with a TP site 10 to 25 amino acid residues upstream. Using this approach, we retrieved a list of 34 novel putative cyclin F substrates (Figure 4F), among which, NEK9 was successfully validated as a cyclin F interactor through IP-WB (Figure S4D), suggesting the TP plus RXIF motif identified in this study can be used to identify uncharacterised cyclin F substrates.

### EXO1 T824 phosphorylation by CDK1/cyclin A is required for interaction with cyclin F and subsequent ubiquitination

As mentioned earlier, T824 in EXO1 closely resembles T33 in RRM2, whose phosphorylation was found to be essential for initiating RRM2-cyclin F interactions ^15^, we therefore wondered whether the cyclin F-EXO1 interaction is also phosphorylation-dependent, and whether the phosphorylation takes place on T824. We first tested the T824 dependence by mutating it to alanine. As presented in Figure 4D and Figure 5A, the T824A mutation abrogated cyclin F-EXO1 interaction. Moreover, unbiased analysis via LC/MS of IP-purified EXO1 WT and EXO1 T824A identified cyclin F as the major differential interacting protein (Figure 5B). And as expected, the EXO1 T824A could not be ubiquitinated by SCF^cyclin^ ^F^ (Figure S4C). On the contrary, when the serine 815 residue was mutated to alanine, there was no detectable difference in EXO1-cyclin F interaction and EXO1 ubiquitination (Figure 4D, Figure 5A, and Figure S4C). These data suggest that EXO1-cyclin F interaction and SCF^cyclin^ ^F^-mediated ubiquitination are both T824-dependent.

**Figure 5.**
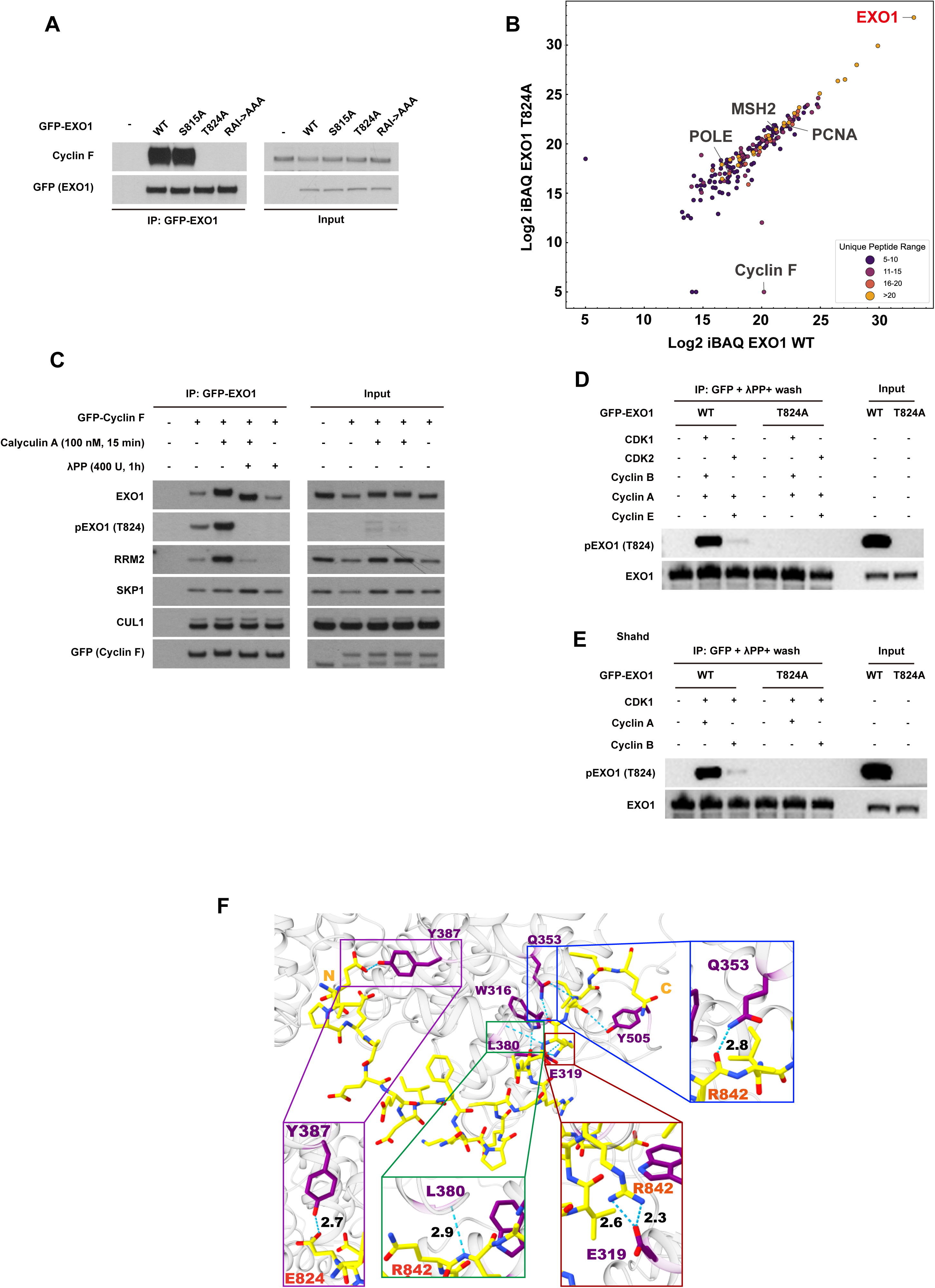
Cyclin A/CDK1 Mediated pT824 is Required for Initiating Cyclin F-EXO1 Interaction. A. Immunoblotting after immunoprecipitation (IP) of GFP-EXO1 WT, GFP-EXO1 S185A, GFP-EXO1 T824A, GFP-EXO1 RAI842AAA mutants. B. Quantification of mass spectrometry results after Flag-immunoprecipitation, comparing Flag-EXO1 WT vs Flag-EXO1 T824A. Unique peptide ranges are labelled with the indicated colours. C. Immunoblotting after immunoprecipitation (IP) of GFP-cyclin F after treatment with Calyculin A or λPP as indicated. λPP treatment was conducted on beads after IP. D. Immunoblotting of *in vitro* phosphorylation assay using GFP-EXO1 WT or GFP-EXO1 T824A purified from HEK293T cells as substrates and the indicated CDK-cyclin combinations as kinases. Immunoprecipitation purified GFP-EXO1 WT or T824 mutant were dephosphorylated on-beads before *in vitro* phosphorylation. E. Immunoblotting of *in vitro* phosphorylation assay using GFP-EXO1 WT or GFP-EXO1 T824A purified from HEK293T cells as substrates and the cyclin A/CDK1 or cyclin B/CDK1 as kinases. Immunoprecipitation purified GFP-EXO1 WT or T824 mutant were dephosphorylated on-beads before *in vitro* phosphorylation. F. Docking of EXO1 peptide to cyclin F protein using COSMIC2. The peptide sequence from L823 to Q846 of EXO1 was used for the docking prediction. To mimic the phosphorylated status of EXO1, T824 was substituted to an E before docking. Cyclin F structure used in the docking is a predicted structure by AlphaFold Protein Structure Database. The iptm+ptm value of 0.7984765224933627.

To establish whether T824 is phosphorylated *in vivo*, we raised antibodies specific to phosphorylated T824 on EXO1 and validated them via IP-WB (Figure S5A) and dot blot (Figure S5B). One out of the two affinity-purified polyclonal antibodies were found to be highly specific to EXO1 pT824. To establish that the antibody specifically recognises the phosphorylated form of EXO1, we treated cells with calyculin A, a potent inhibitor of protein phosphatases 1 (PP1) and 2A (PP2A), to elevate global phosphorylation in cells. In addition, we also treated immunoprecipitated proteins with λ phosphatase (λPP), a Mn^2+^-dependent protein phosphatase with activity towards phosphorylated serine, threonine, and tyrosine. Treatment with calyculin A was found to elevate the EXO1 pT824 signal, while treatment with λPP completely removed it (Figure 5C), indicating EXO1 is phosphorylated on T824 *in vivo*. Moreover, calyculin A induced hyperphosphorylation also promoted cyclin F’s interaction not only with the classic cyclin F substrate RRM2, but also with EXO1. However, λPP treatment after calycuclin A only brought cyclin F-RRM2 interaction back to the untreated level and had no effect on the cyclin F-EXO1 interaction (Figure 5C). Taken together, these data suggest that T824 phosphorylation is indeed required to initiate cyclin F-EXO1 interaction but is dispensable for the maintenance of the interaction.

To identify the kinase responsible for T824 phosphorylation, we used the home-made site-specific phosphorylation antibody to screen a panel of inhibitors targeting kinases involved in the cell cycle and DNA damage responses. While several inhibitors impacted cyclin F levels and its interaction with EXO1, only RO-3306 (CDK1 inhibitor) and K03861 (CDK2 inhibitor) induced a notable reduction in both cyclin F-EXO1 interaction and EXO1 T824 phosphorylation (Figure S5C). BI6727 (PLK1 inhibitor) and CKIIi VIII (Casein Kinase II inhibitor) also drastically downregulated cyclin F-EXO1 interaction, likely due to the reduction in endogenous cyclin F protein levels as described previously ^33^, but they did not impact on EXO1 T824 phosphorylation (Figure S5C). Thus, the kinases phosphorylating EXO1 on T824 are likely to be CDKs in line with the degradation of EXO1 at G2 and M when CDK activity peaks.

To further define the specific cyclin/CDK partner(s) responsible for the T824 phosphorylation, we implemented an *in vitro* phosphorylation assay using CDK2/cyclin E, CDK2/cyclin A, CDK1/cyclin A, and CDK1/cyclin B. A significant increase in EXO1 T824 phosphorylation was detected only upon the addition of CDK1 into the reaction but not CDK2, regardless of the types of the cyclin proteins (Figure 5D). Moreover, as an effort to further identify the responsible cyclin protein, CDK1/cyclin A and CDK1/cyclin B were tested for their capability in phosphorylating EXO1 at T824 and only CDK1/cyclin A significantly increased EXO1 T824 phosphorylation (Figure 5E). Taken together, these results demonstrate that cyclin A/CDK1 phosphorylates EXO1 at T824.

Both RRM2 and EXO1 rely on the threonine phosphorylation step and RxIF motif for cyclin F interaction. The mechanism of interaction can be explained by the phosphorylation being a priming mechanism that triggers exposure of the RxIF motif, or by the phosphorylation being an additional binding site that directly interacts with cyclin F cooperatively with the RxIF motif. The latter possibility was tested using Alphafold-Multimer to dock cyclin F with the EXO1 peptide spanning from the phosphorylation site to the end of the RxIF motif (Figure 5F). Interestingly, the model highlights hydrogen bonds between EXO1’s R842 residue and cyclin F’s E319 residue, which closely resemble the interaction between p27’s R30 residue and cyclin A’s E220 residue ^34^. In addition, the same model predicts a hydrogen bond forming between EXO1’s T824 (mutated to E824 to mimic the phosphorylated residue) and cyclin F’s Y387, indicating a potential second valence mediating the cyclin F-EXO1 interaction. The results above suggest a multivalent binding mechanism similar to that used by other ligases ^35^ and they outline a possible multi-step procedure to prevent uncontrolled degradation of CDK/cyclin substrates by SCF^cyclin^ ^F^ at the G2 and M transition.

### EXO1 R842A mutation phenocopies *CCNF* K/O

As shown in Figure 3A, EXO1 is the major mediator of IR sensitivity upon *CCNF* loss, and downregulating the stabilised EXO1 can reverse *CCNF* K/O’s radiosensitisation effect (Figure 3C and Figure 3D). It is reasonable to hypothesize that EXO1 stabilisation should phenocopy *CCNF* loss in regard to IR sensitivity. To test this hypothesis in a refined system, we generated cell lines that, upon doxycycline treatment, express HA-tagged wild-type EXO1 (WT) or EXO1 R842A (which, according to previously demonstrated data, abrogates cyclin F-EXO1 interaction and EXO1 ubiquitination). As validation in Figure 6A shows, both WT and R842A mutant EXO1 were successfully expressed upon induction with doxycycline, with R842A demonstrating a higher protein level in the input (Figure 6A). IP-WB also confirmed that the R842A mutation significantly reduced its interaction with cyclin F (Figure 6A). Moreover, CHX chase experiments also showed a significantly longer protein half-life of the R842A mutant, proving that EXO1 R842A is indeed resistant to SCF^cyclin^ ^F^-mediated degradation (Figure 6B). In a cell cycle synchronisation experiment, we also observed that EXO1 WT was degraded in G2 and M when the level of cyclin F peaks, while EXO1 R842A was not (Figure 6C), confirming that cell cycle regulation of EXO1 is directly mediated by cyclin F through the interaction with the RxIF motif. Lastly, enrichment of endogenous ubiquitinated proteins using UBE pulldown assay confirmed that EXO1 WT can be ubiquitinated while EXO1 R842A cannot (Figure 6D), validating a crucial role of the R842 in potentiating EXO1 ubiquitination.

**Figure 6.**
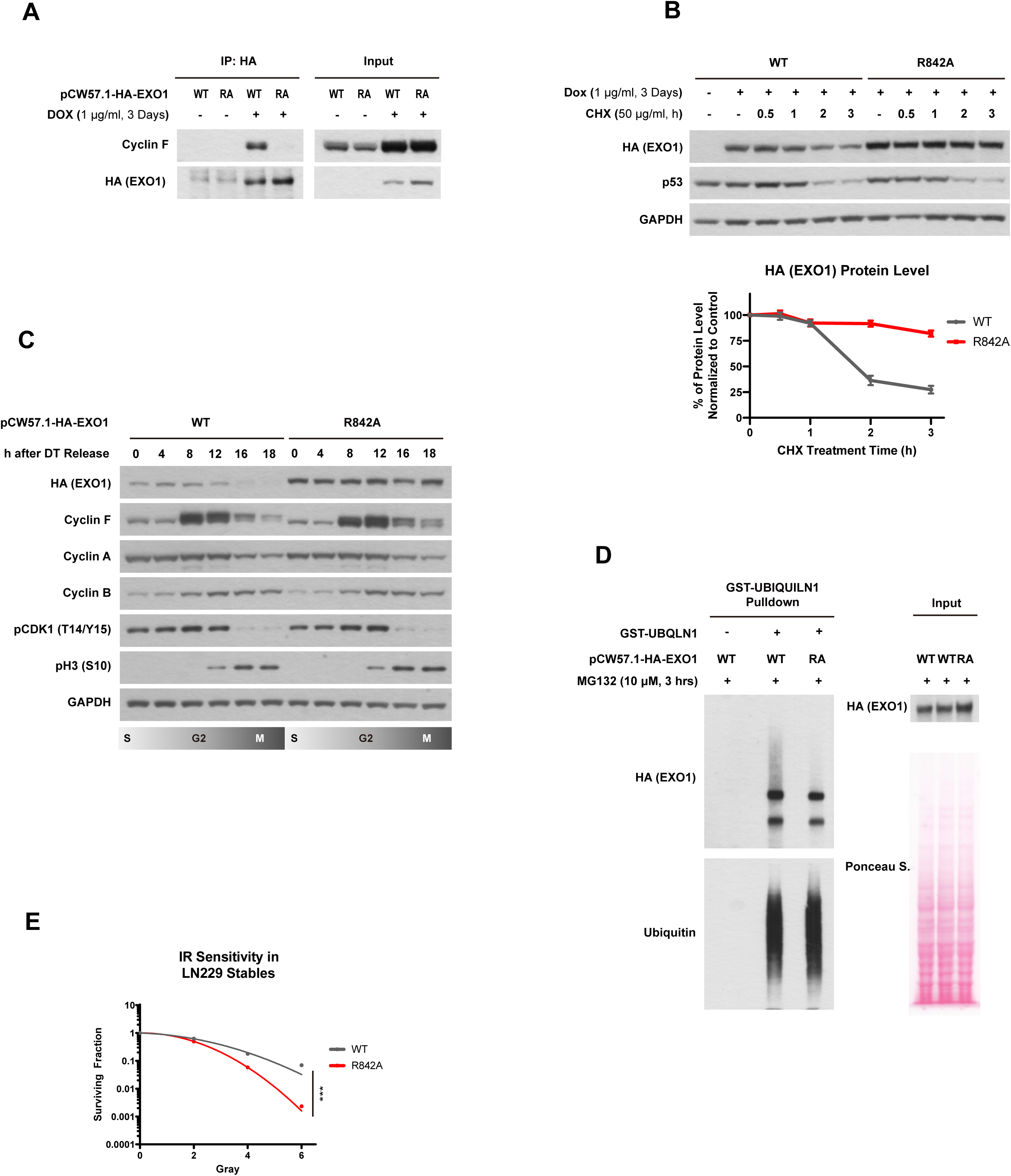
EXO1 R842A mutation phenocopies *CCNF* K/O. A. Immunoblotting after immunoprecipitation (IP) of stably integrated HA-EXO1 WT or HA-EXO1 R842A in LN229 using a doxycycline inducible promoter. Cells were treated with 1 μg/ml doxycycline for 3 days to induce expression. B. Immunoblotting after treating cells described in A with Cycloheximide (CHX) for the indicated time -*upper panel*. Relative quantification of EXO1 protein levels in HA-EXO1 WT or HA-EXO1 R842A after normalisation with EXO1 levels at T0 for each cell line - *bottom panel*. Error bars: standard deviations of three biological replicates; C. Immunoblotting of cell cycle synchronised cells described in A *via* double thymidine (DT) block-release. D. Immunoblotting after isolation of endogenous ubiquitinated proteins using Ubiquitin binding entity (UBE) in cells described in A. E. LN229 expressing HA-EXO1 WT or HA-EXO1 R842A were seeded for colony formation assay and challenged with the indicated dose of IR. 7 days after IR, cells were stained with crystal violet and counted. Error bars represent standard deviations of three biological replicates. Statistical analysis two-tailed unpaired t test. ** indicates P ⩽ 0.01, *** indicates P ⩽ 0.001, and **** indicates P ⩽ 0.0001.

Given that IR sensitivity in cyclin F-depleted cells can be fully rescued by concomitant EXO1 depletion (Figure 3A), EXO1 R842A is expected to recapitulate the radiosensitisation phenotype observed in *CCNF* K/O cells (as shown in Figure 1D). Indeed, the colony formation assay in Figure 6E demonstrated that cells expressing the EXO1 R842A mutation are significantly more sensitive to IR compared to cells expressing EXO1 WT. Moreover, the steady-state level of γH2Ax was also increased in R842A mutant (Figure S6C), further confirming the radiosensitisation effect of EXO1 stabilisation.

Interestingly, a point mutation at R842 was identified in a glioblastoma patient by the Cancer Genome Atlas Program (TCGA) (Figure S6A). Although the arginine residue was mutated to isoleucine in this case, the R842I mutation also disrupts cyclin F-EXO1 interactions (Figure S6B). It is tantalising to speculate that the good prognosis of this patients at 6 months, might be due to an exceptional response to radiotherapy as observed in other cases bearing alterations in other DDR genes ^36^. Unfortunately, the patient was not followed beyond 6 months when recurrence occurs, but this observation suggests nonetheless that the cyclin F-EXO1 axis can indeed be dysregulated in GBM via EXO1 mutation at the degron. It is possible that these and other mutations impacting IR-related pathways would be more evident in tumours that recur after radiotherapy, but this cannot be confirmed as repeat surgery is rare, limiting sample availability.

Altogether, the effect of *CCNF* K/O on EXO1 stability in G2 and M was successfully recapitulated by a single point mutation in EXO1 at R842.

### EXO1 R842A mutation leads to hyper-resection and chromosome aberrations after IR

As demonstrated thus far, the regulation of EXO1 is mainly driven by cyclin F during the cell cycle, and dysregulation of EXO1 upon either cyclin F loss or EXO1 R842A mutation leads to IR hypersensitivity. However, the mechanism by which aberrantly stabilised EXO1 leads to IR hypersensitivity remains to be elucidated.

Since EXO1 is crucial for DNA long-range end resection, a process required by homologous recombination repair (HR) to generate ssDNA at DSBs, we started the mechanistic study by measuring the formation of ssDNA. As shown in Figure S7A, a significant increase in the percentage of cells with ssDNA was observed upon *CCNF* K/O in HeLa (Figure S7A), which was rescued by concomitant knockout of *EXO1* (Figure S7A), indicating the elevated resection is indeed EXO1-mediated. Similarly, more cells with ssDNA foci were detected in cells expressing EXO1 R842A (Figure 7A). As more cells undergoing resection does not necessarily mean longer resection tracks, we also assessed DNA end resection directly at the level of individual DNA molecules via SMART assay (Single Molecule Analysis of Resection Tracks assay) and observed a significant increase in the length of resection tracks in the R842A mutant (Figure 7B). Taken together, these results suggest that *CCNF* K/O or EXO1 R842A mutation-mediated EXO1 stabilisation can lead to increase in both prevalence and length of DNA resection— a phenomenon known as DNA hyper-resection.

**Figure 7.**
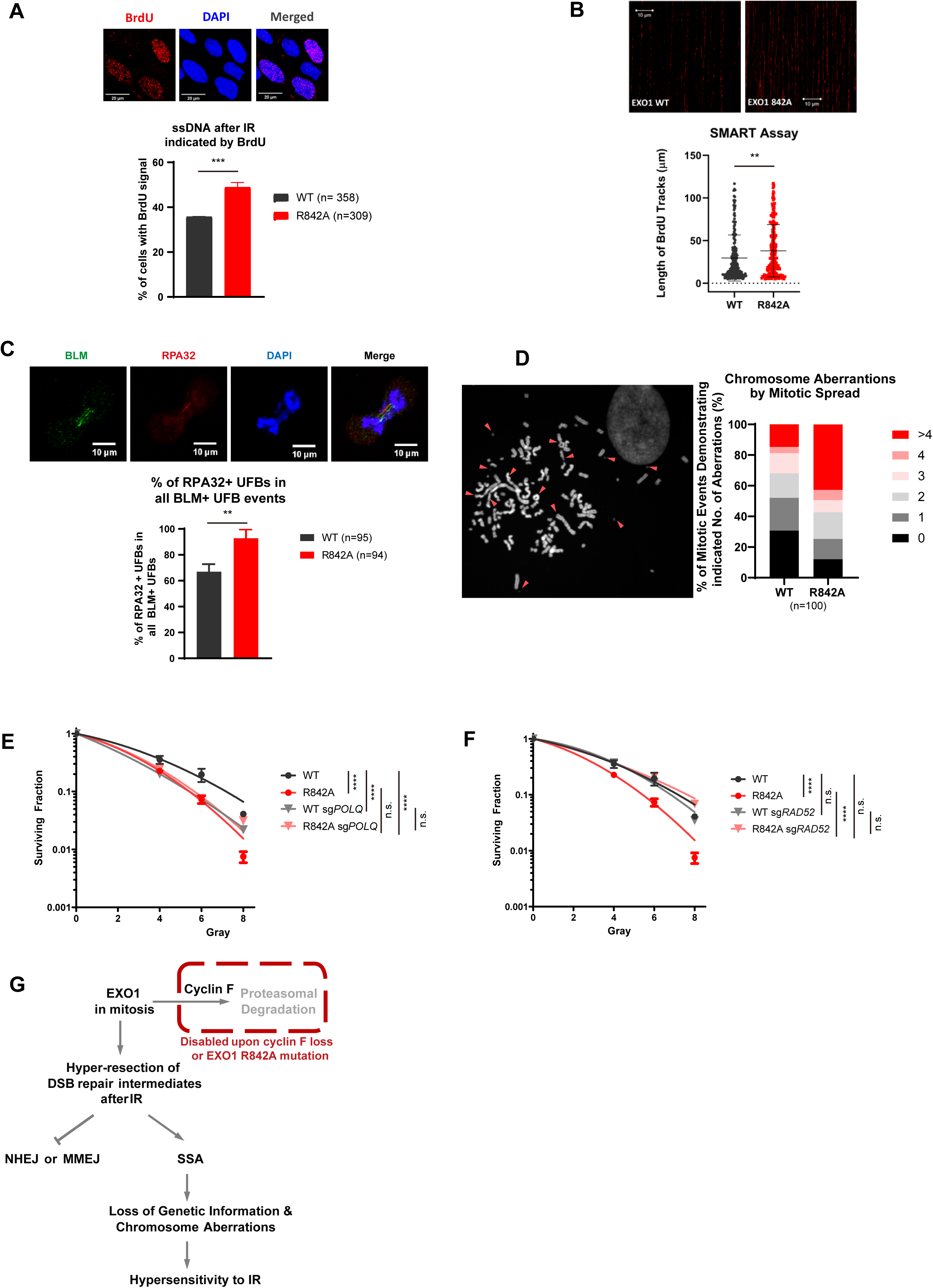
EXO1 R842A Mutant Leads to Hyper-Resection and Toxic Single Strand Annealing after IR. A. Representative images of cells containing BrdU signal in LN229 expressing HA-EXO1 or EXO1 R842A mutant. After doxycycline induction for 3 days, cells were re-seeded for 50% density and labelled with 10 μM BrdU for 24 hours before being treated with 10 Gy IR and allowed to recover for 3 hours. Quantification of the signal retrieved in the indicated cells (n). Statistical analysis using two-tailed unpaired t test. *** indicates P ⩽ 0.001. B. Representative DNA fibers using Single Molecule Analysis of Resection Tracks (SMART) assay in LN229 expressing HA-EXO1 or EXO1 R842A mutant. Quantification of fiber length in 300 events. Statistical analysis using Mann-Whitney U test. ** P =0.0059, U=17873.5, Z=-2.7533. C. Quantification of RPA32 positive anaphase ultra-fine bridges (UFBs) in LN229 expressing HA-EXO1 or EXO1 R842A mutant. After doxycycline induction for 3 days, cells were treated with 10 Gy IR and allowed to recover for 12 hours before subjected to UFB detection. Representative images – *upper panel*. Error bars represents standard deviation of three biological replicates. n represents total events quantified in three replicates. Statistical analysis was done using two-tailed unpaired t test. ** indicates P ⩽ 0.01. D. Quantification of chromosome alterations in RPE-1 expressing HA-EXO1 or EXO1 R842A mutant after mitotic spreading. After doxycycline induction for 3 days, hTERT RPE-1 cells with pCW57.1-EXO1 WT or R842A were treated with 10 Gy IR and allowed to recover for 72 hours before further treated with colcemid for 3 hours to enrich metaphase cells. Cells were then harvested for mitotic spreading. A total of 100 metaphase events for each condition were analysed. Percentages of mitotic events demonstrating 0, 1, 2, 3, 4, or >4 aberrations were calculated and plotted. E. LN229 expressing HA-EXO1 WT or HA-EXO1 R842A and transfected with a control sgRNA or sgRNA targeting *POLQ* as indicated were seeded for colony formation assay and challenged with the indicated dose of IR. 14 days after IR, cells were stained with crystal violet and counted. Error bars represent standard deviations of three biological replicates. F. LN229 expressing HA-EXO1 WT or HA-EXO1 R842A and transfected with a control sgRNA or sgRNA targeting *RAD52* as indicated were seeded for colony formation assay and challenged with the indicated dose of IR. 14 days after IR, cells were stained with crystal violet and counted. Error bars represent standard deviations of three biological replicates. G. Schematic representation of cyclin F-EXO1 axis in the control of DSB repair. EXO1 accumulation prevents MMEJ in mitosis but promotes uncontrolled SSA, which ultimately hyper-sensitizes cells to DSBs induced by IR treatment.

Since the effects of cyclin F loss on EXO1 stability are seen in mitosis, we sought further evidence of extensive resection in this phase through probing for single-stranded anaphase ultra-fine bridges (UFBs) characterised by replication protein A2 (RPA32) coating. UFBs are also identifiable by the localisation of Bloom’s syndrome helicase (BLM) and PLK1-interacting checkpoint helicase (PITCH) to the ssDNA in unresolved recombination intermediates or at common fragile sites. UFBs coated with RPA32 represent stretches of unrepaired ssDNA which could eventually fracture at the end of mitosis ^37, 38^. Examples of RPA32-positive and RPA32-negative UFBs are shown in the upper panel of Figure 7C. Quantification of these events in cells expressing EXO1 WT or EXO1 R842A showed a significantly higher incidence of RPA32-positive UFBs in mutant cells (Figure 7C lower panel). Similarly, UFBs were increased in HeLa cells with *CCNF* K/O or *EXO1* K/O (Figure S7B) and this was rescued by concomitant *EXO1* K/O upon *CCNF* K/O background. The experiments above further supported the idea that both *CCNF* K/O and EXO1 R842A mutation lead to DNA hyper-resection.

DNA Hyper-resection creates long strand of mechanically fragile sites in the genome, the fracture of which could eventually lead to chromosome breakage during segregation. Moreover, hyper-resection may also expose extensive single-stranded homologies among random genomic regions, which could facilitate unwanted HR among different chromosomes or within the same chromosome, causing chromosome fusion or the formation of radial chromosomes, which, in turn, increases the chance of asymmetric chromosome division and aneuploidy. This speculation is indeed supported by the significantly higher incidence of chromosome aberrations observed after IR in hTERT RPE-1 cells expressing EXO1 R842A (Figure 7D). As expected, the majority of these aberrations were radial chromosomes, chromosome fusions, chromosome breakage, and aneuploidy.

The presence of extensive DNA resection is a sign of uncontrolled EXO1 activity at DSBs. As extensive resection can prevent the efficient execution of multiple DSB repair pathways ^39, 40^, we decided to identify DSB repair pathways that become defective upon aberrant EXO1 stabilisation. To this end, we concomitantly genetically ablated HR, NHEJ, SSA, or MMEJ in cells expressing EXO1 WT or EXO1 R842A and assessed IR sensitivity. After removing *XRCC4* (essential for NHEJ), *POLQ* (necessary for MMEJ), *RAD51* (important for HR), and *RAD52* (required for SSA), deficiencies in NHEJ or HR were found to have an additive effect on the radiosensitisation phenotype elicited by EXO1 R842A expression. In other words, removal of *XRCC4* or *RAD51* promotes IR sensitivity independently of EXO1 stabilisation (Figure S7C and S7D). In contrast, the effect of removing *POLQ* was epistatic to EXO1 R842A expression, indicating that MMEJ is compromised in cells with high levels of EXO1 R842A (Figure 7E). Our findings show that stabilised EXO1 in mitosis counteracts POLQ-mediated MMEJ, in line with recent findings showing that MMEJ operates mostly in mitosis ^41^. Our findings show that EXO1 must be removed in mitosis to allow MMEJ, potentially by preventing removal a POLQ template or generation of extensive resection tracts that POLQ is unable to fill. Interestingly, genetic ablation of *RAD52* in the EXO1 R842A cell line fully rescued the IR sensitivity promoted by EXO1 stabilisation (Figure 7F), showing that EXO1 stabilisation-induced hyper-resection leads to DNA damage and cell death through SSA. It is possible that cells attempt SSA to resolve the DSBs left in mitosis, but SSA in these conditions is deleterious as it promotes chromosomal losses and rearrangements inducing cell death.

## Discussion

To identify novel determinants of IR sensitivity, we performed a focused and high-resolution CRISPR screen in radioresistant GBM cell lines. The screen identified both well-established and new factors modulating IR sensitivity. Among the latter, *CCNF* depletion was validated to cause radiosensitisation. Subsequent investigation revealed that SCF^cyclin^ ^F^ facilitates EXO1 ubiquitination and the subsequent proteasomal degradation during G2 and M cell cycle phases. EXO1 accumulation upon cyclin F loss accounts for the IR hypersensitivity observed in *CCNF* K/O cells. EXO1 was previously identified as a substrate of SCF^cyclin^ ^F^ ^9, 11^ following DNA damage. Our study instead revealed a cell cycle-dependent regulation mechanism. Moreover, we showed that the cyclin F-EXO1 interaction requires CDK1/cyclin A -mediated phosphorylation at T824 as a priming signal, and that the interaction is mediated by the cyclin domain on cyclin F and the RAIF motif at the C-terminus of EXO1. Importantly, expression of the non-degradable R842A EXO1 mutant recapitulated the IR hypersensitivity phenotype observed in *CCNF* K/O conditions. Further investigation suggests that the presence of EXO1 in G2 and M promotes resection, preventing the repair by MMEJ. It is likely that cells attempt to repair these DSBs using SSA instead. In these circumstances, upon DNA DSB induction, the uncontrolled SSA leads to chromosomes fusions and breakage, which ultimately induces cell death.

UBA3 inhibitor MLN4924 was previously reported to increase IR sensitivity in cells and *in vivo* ^42^. However, the individual CRL adaptors and the mechanisms responsible for such hypersensitivity remained unclear. Our screen identified multiple components of SCF complexes, including but not limited to genes encoding CUL1 and cyclin F, which opens several routes of investigation to explain the pleiotropic effects of MLN4924.

As aberrant stabilization of mutant EXO1 in mitosis leads to DNA hyper-resection, the product of which is known to be a poor substrate for MMEJ, our findings on cyclin F-mediated EXO1 degradation in G2 and M explain why MMEJ is mainly restricted to mitosis, providing a novel insight into the mechanism of pathway choice between SSA and MMEJ.

A key mediator of MMEJ is the DNA polymerase POLQ. Several POLQ inhibitors are being developed and are entering clinical trials as they demonstrate exciting anti-cancer potential ^43^. Given that *CCNF* depletion- or EXO1 R842A mutation-mediated hyper-resection leads to defective MMEJ, exploiting the cyclin F-EXO1 pathway could be an alternative approach to POLQ inhibition to restrict MMEJ and, therefore, to sensitise cells to IR. Moreover, similar to inhibiting POLQ, cyclin F inhibition or EXO1 stabilisation also induces the formation of micronuclei (Figure S7E) ^16^, which is known to facilitate inflammatory signaling activation after IR and to synergize with radiotherapy ^44^. It is therefore reasonable to believe cyclin F is a promising target for increasing the efficacy of radiotherapy in glioblastoma and other tumour types.

We showed that the degradation of EXO1 during G2 and M is initiated by CDK1/ cyclin A-mediated phosphorylation and is cell cycle-dependent. We and others ^9, 11^ also observed (Figure 3D and S3A) that SCF^cyclin^ ^F^ mediates EXO1 degradation upon IR, UV, and other genotoxic stimuli. Therefore, it is reasonable that additional regulatory phosphorylation events on EXO1 could exist and are required for DNA damage-induced EXO1 degradation.

Our study also focused on the analogy between degrons found in RRM2 and EXO1, and uncovered a recognition site specific to cyclin F and distinct from the classical Cy motifs (RxL or KxL). Our investigation reveals that cyclin F specifically recognises the RxIF motif, which we name F-deg, potentially resolving the conundrum of how cyclin F achieves substrate specificity instead of degrading all proteins containing Cy motifs. It is worth mentioning that our model does not exclude the possibility that cyclin F is still able to interact with the canonical Cy motif, but defines how cyclin F specifically distinguishes substrate interactors from non-substrate interactors.

In addition to helping elucidate a pathway for the repair of DSBs after IR, our CRISPR screen highlighted many novel factors with unknown roles. Although the use of immortalized cell lines in this screen prevents true recapitulation of transcriptional features in human glioblastoma cells ^45^, these factors represent potential weaknesses of glioblastoma cells, exploitable in targeted therapies. Further expansion of the screen in glioma stem cells or in *in vivo* models will validate the work presented here in a more clinically relevant context.

## Material and Methods

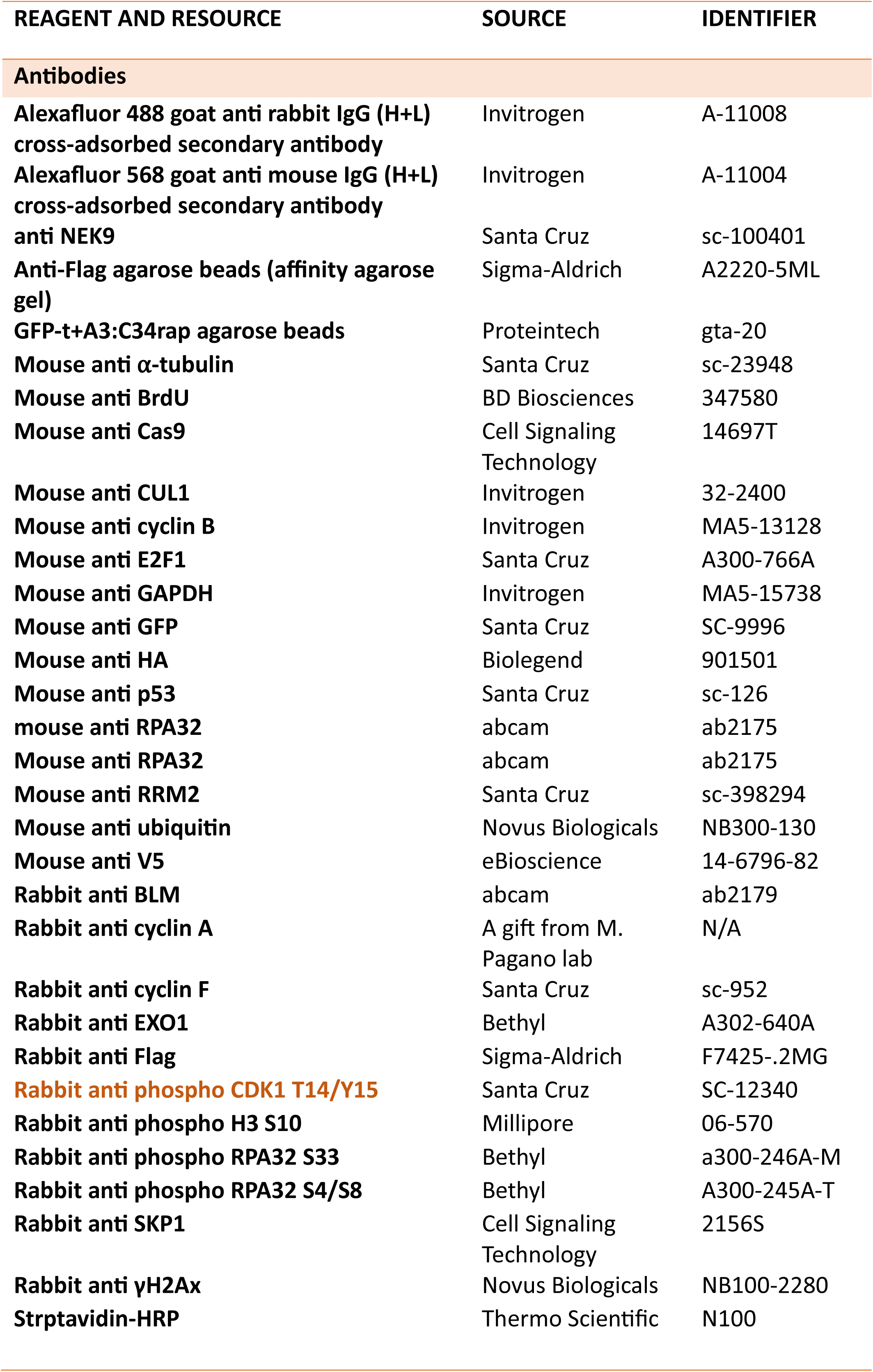
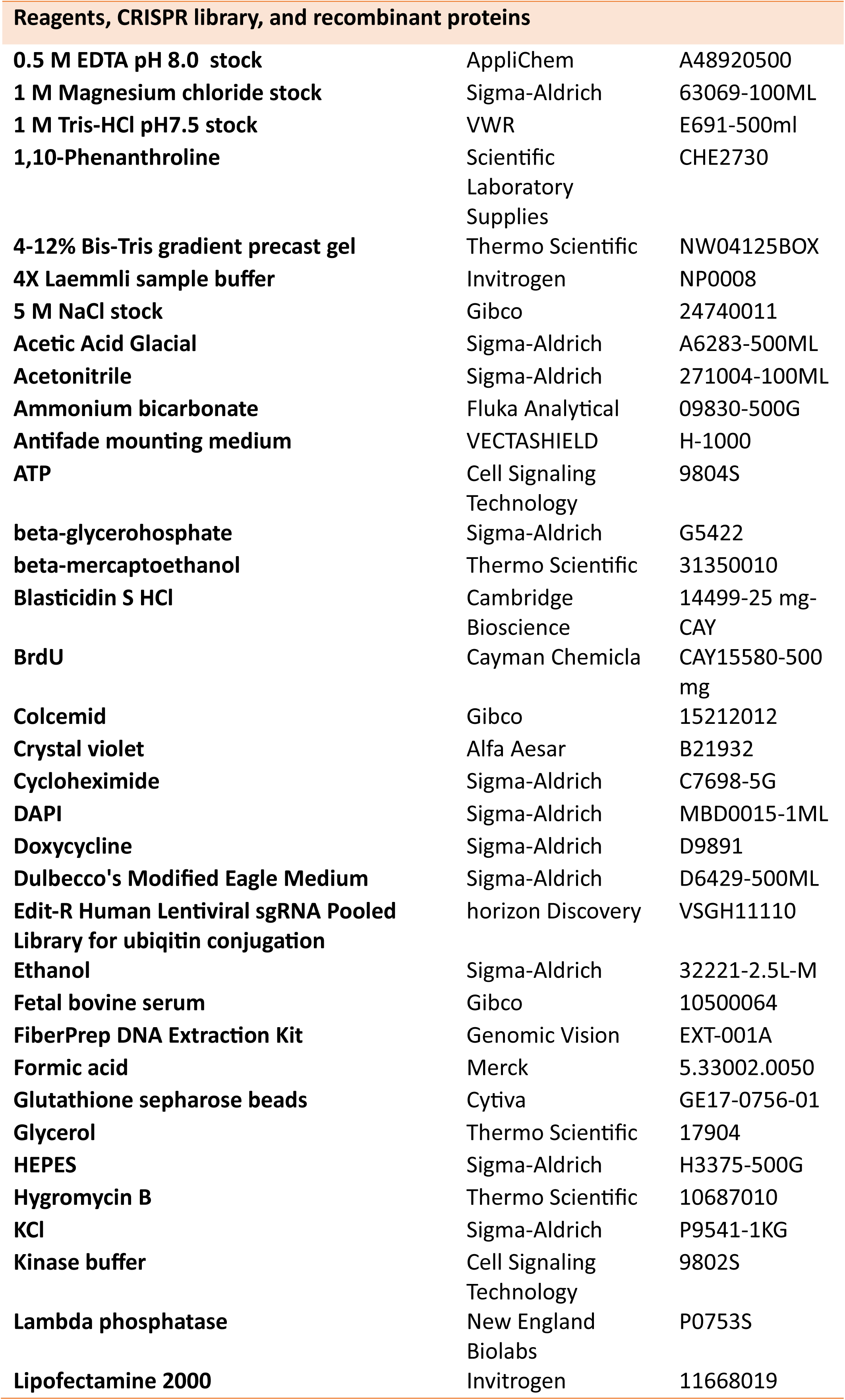
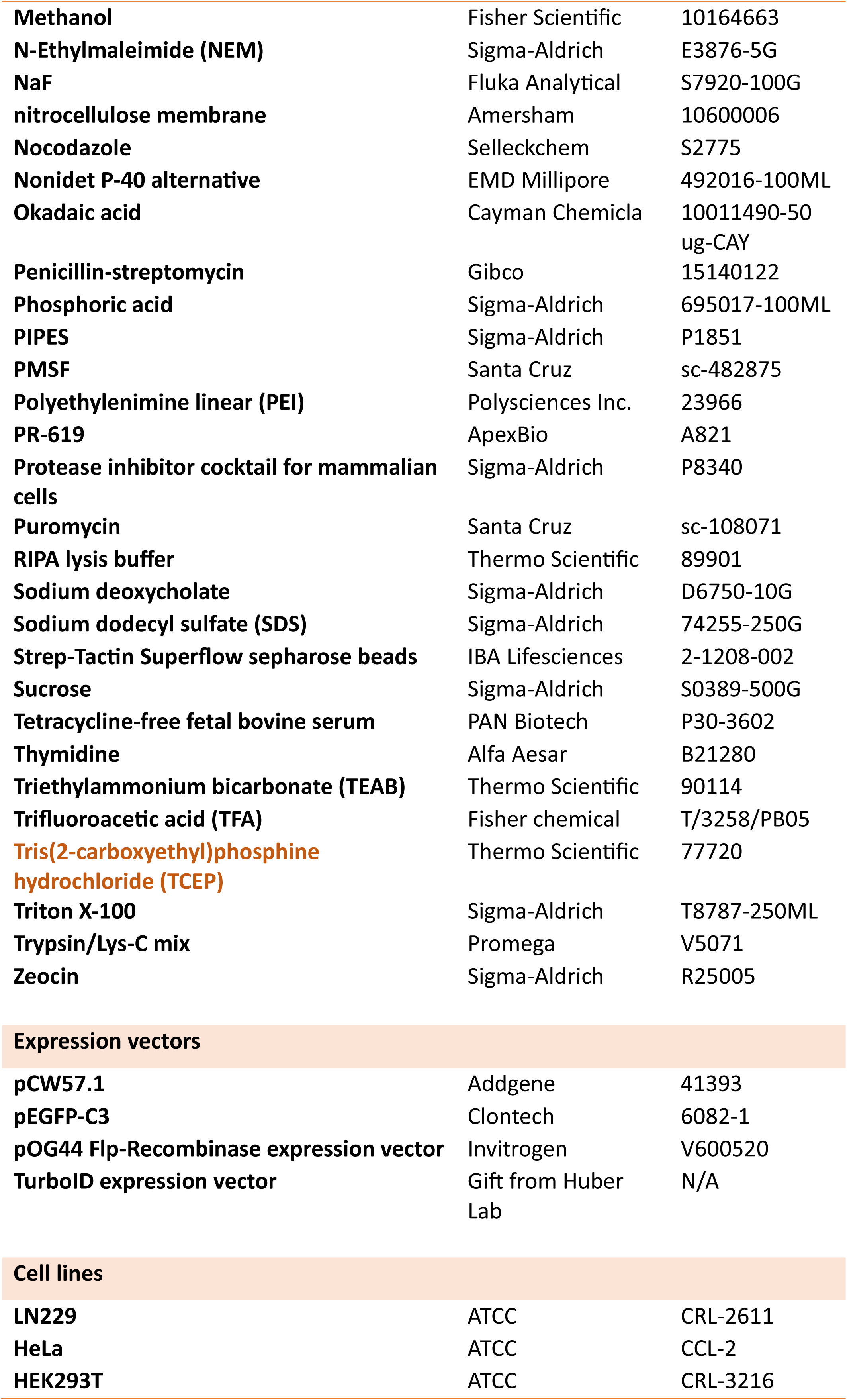

### CRISPR Screen and Data Analysis

Viral particles for the Edit-R Human Lentiviral sgRNA Pooled Library - UB conjugation (cat. n. VSGH11110) were purchased from Dharmacon. Viral particles were transduced into an LN229 Cas9-stable line with a multiplicity of infection (MOI) of 0.3 (less than 3% of all cells will be infected by more than one viral particle). We performed the transduction to ensure a fold of representation of 750 (meaning that, on average, the each guide was transduced into 750 different cells in the pooled population) and kept this fold of representation throughout the screen. Each sample was run in biological quadruplicates. After treatment, genomic DNA (gDNA) was extracted using Qiagen DNA extraction kit (Cat. No. / ID: 69504) according to manufacturer’s instructions. sgRNA sequences harboured were amplified by PCR using Edit-R Pooled sgRNA Indexing PCR and Sequencing Primer Kit A (PRM10184) and Edit-R Pooled sgRNA Indexing PCR and Sequencing Primer Kit B (PRM10185). PCR product were sequenced on a Hiseq4000 at the Welcome Trust center for Human Genetics, Oxford. The sequencing data was converted into readcounts and statistical analyses were run using CRISPRanalyzeR (http://crispr-analyzer.dkfz.de).

### Over-representation analysis (ORA)

ORA was performed using the EnrichR program ^46^. In short, statistically significant genes conferring either IR sensitivity or resistance ([Z-Ratio]>1.96) were combined as one gene set and submitted for analysis on the EnrichR website. The dataset was ranked by Kappa score ^47^. Pathways with a Kappa score of ≥ 0.65 were considered high-confidence hits. Pathways with a Kappa score of ≥ 0.5 but < 0.65 were considered confident identifications, and those with a Kappa score of ≥ 0.35 but < 0.5 were considered possible hits.

### Cell Culture

LN229, HEK293T, H4, SW1088, DBTRG, A172 and HeLa cells were obtained from ATCC. All cell lines were cultured in DMEM (Sigma, D6429-500ML) containing 10% fetal bovine serum (FBS) (Life, 10500064) and the mixture of 100U/ml penicillin and 100 ug/ml streptomycin (Life, 15140122). HeLa cells with *CCNF* and/or *EXO1* knocked out were generated using a CRISPR system (sgRNA sequences: Table above). LN229 cells transduced with pCW57.1 lentiviral expression system were maintained in tetracycline-free media (DMEM plus 10% tetracycline-free FBS (PAN Biotech, P30-3602)). Expression of pCW57.1-EXO1 was induced by culturing the EXO1-stable cells in media containing doxycycline (Sigma-Aldrich, D9891, at a final concentration of 1 μg/ml) for 3 days before seeding for experiments. 1 μg/ml doxycycline was present at all times and was replenished every 3 days during experiments unless otherwise specified.

### Colony formation Assay

Cells were counted and seeded at 400 cells per well in 6-well plates and cultured for 6 hours before being challenged with the indicated IR doses using a closed source gamma irradiator. After IR treatment, cells were allowed to propagate for 7 days (for HeLa derived cell lines) or 14 days (for LN229 derived cell lines). Colonies were fixed and visualised using crystal violet solution (50% methanol, 10% ethanol, 0.3% crystal violet) for 10 minutes at room temperature with gentle agitation (10-20 rpm). After rinsing with water and airdrying, colonies were counted using GelCount™ mammalian-cell colony counter (Oxford OPTRONIX). All colony formation assays presented in this study have been repeated at least three times. And error bars represent standard deviation of three biological replicates. Statistical analysis was done using two-tailed unpaired t-tests.

### Generation of TurboID-Cyclin F Stable Cell Line

Flp-In™ T-REx™ HEK293 cells (Invitrogen, R78007) containing a single genomic FRT site and stably expressing the Tet repressor were cultured in DMEM medium supplemented with 10% FBS, 100 μg/ml zeocin, and 15 μg/ml blasticidin. The medium was replaced with fresh medium containing no antibiotics before transfection. For cell line generation, Flp-In HEK293 cells were co-transfected with the pCDNA3-TurboID-cyclin F plasmid and the pOG44 Flp-Recombinase Expression Vector (Invitrogen, V600520) for co-expression of the Flp-recombinase using Lipofectamine 2000 transfection reagent (Invitrogen, 11668019). Two days after the transfection, cells were selected in hygromycin-containing medium (100 μg/ml) for 2–3 weeks. To validate TurboID-cyclin F expression, cells were cultured in media containing 1.3 μg/ml doxycycline for 24 h to induce TurboID-Cyclin F expression before levels were checked by immunoblot.

### Mass Spectrometry (MS) Sample Preparation for TurboID-Cyclin F pulldown

When TurboID-cyclin F Flp-In™ T-REx™ HEK293 cells reached 80% confluency in 15 cm dishes, 1.3 μg/ml doxycycline was added for 24 h to induce the expression of TurboID-cyclin F. Cells were further incubated with 50 µM biotin for 3 h to label proteins that came into close proximity with TurboID-cyclin F in cells. Cells were harvested by scraping and washed 3 times with PBS. For streptavidin pulldown of all biotin-labelled proteins (potential cyclin F interactors), cell pellets were thoroughly resuspended in 1 ml of RIPA buffer (50 mM Tris-HCl pH 8.0, 150 mM NaCl, 1% Triton, 1 mM EDTA, 0.1% SDS with Protease Inhibitor cocktail (Sigma-Aldrich, P8340) and incubated on ice for 15 min. Insoluble material was removed by centrifugation. Cleared lysates were then incubated on a rotating wheel at 4°C with 50 μl pre-equilibrated Strep-Tactin® Superflow Sepharose beads (IBA, 2-1208-002) for 1 h. The suspension was then loaded on a Mini Bio-Spin® Columns (Bio-Rad, 732-6207) to collect the beads. The beads were washed two times with 1 mL RIPA buffer, three times with HNN buffer (50 mM HEPES pH 7.5, 150 mM NaCl, 50 mM NaF), and two times with 100 mM NH_4_HCO_3_ solution before being transferred to 2 ml Eppendorf tubes in 400 μl NH_4_HCO_3_ solution. For proteolysis, the sample was centrifuged at 200 x g for 1 min to remove supernatant. Beads were resuspended in 100 μl of 8M Urea in 100 mM NH_4_HCO_3_ solution and incubated at 20 °C for 20 min. Cysteine bonds were reduced with a final concentration of 5 mM Tris(2-carboxyethyl) phosphine hydrochloride (TCEP) for 30 min at 37°C and alkylated in a final concentration of 10 mM iodoacetamide (IAA) for 30 min at room temperature in the dark. Beads were then proteolyzed with Trypsin/Lys-C Mix (Promega, V5071) at a 25:1 protein: protease ratio (w/w) for 4 h at 37°C on an orbital shaker. Urea concentration was then reduced to 1 M via adding 100 mM NH_4_HCO_3_ solution to the sample. Samples were digested overnight at 37°C on an orbital shaker. To remove trypsin, samples were loaded onto C18 spin columns (Thermo Scientific, 89870), washed according to the manual provided by the manufacturer, and eluted with 0.1% TFA (Trifluoroacetic Acid) and 65% acetonitrile. Peptides were then dried in a SpeedVac vacuum concentrator and resuspended in 0.1% TFA and 2% acetonitrile in mass spectrometry (MS) grade water for MS analysis.

### LC-MS/MS Sample Preparation and Data Analysis

Please refer to the section above for MS sample preparation of the proximity labelling TurboID-cyclin F samples. For the identification of the GFP-EXO1 WT versus T824A differential interactome, over-expressed GFP-EXO1 WT and T824A were immunoprecipitated from HEK293T respectively and eluted using 2% SDS buffer. Eluents were then digested using S-Trap micro spin columns following manufacturer’s protocol (Profiti, C02-micro-10). In brief, samples were reduced with 20 mM DTT and alkylated with 40 mM iodoacetamide (30 min each at room temperature in the dark). Samples were then acidified with phosphoric acid (1.2% final concentration) and mixed with binding buffer (100mM triethylammonium bicarbonate (TEAB) in 90% methanol) to a 1:7 ratio (sample: binding buffer). Samples were then transferred onto the S-trap spin column. Proteins in the samples were adsorbed onto columns via centrifugation before being washed with binding buffer five times. 20 μl of 50 mM TEAB containing 1 μg of trypsin was then added to the column and incubated overnight at 37°C. Peptides were eluted with 50 mM TEAB and 2% formic acid in 50% acetonitrile solution and dried using a SpeedVac vacuum concentrator. Before MS, samples were resuspended in 0.1% TFA and 2% acetonitrile in MS grade water.

For both TurboID-cyclin F and GFP-EXO1 MS, samples were analysed by reverse-phase chromatography using an UltiMate 3000 UHPLC connected to an Orbitrap Fusion Lumos (Thermo Fisher). Peptides were trapped on a PepMap C18 trap column (300 µm x 5 mm, 5 µm particle size, Thermo Fisher) and separated on a 50 cm Easy-Spray column (ES803, Thermo Fisher) using a 60-minute linear gradient from 2% to 35% buffer B (buffer A: 5% DMSO, 0.1% formic acid; buffer B: 5% DMSO, 0.1% formic acid in acetonitrile) at a flow rate of 250 nl/min. Eluted peptides were then analysed in the Orbitrap Fusion Lumos in data-dependent mode (DDA) with the advance peak detection (APD) switched on. Full scans were acquired in the Orbitrap at 120k resolution over an m/z range of 400 −1500, AGC target of 4e5, and S-lens RF of 30. MS2 scans were obtained in the Ion trap (rapid scan mode) with a Quad isolation window of 1.6, 40% AGC target and a maximum injection time of 35 ms, with HCD activation and 28% collision energy.

The raw mass spectrometry data were analysed using MaxQuant (v1.6.14). Briefly, files were searched against the UniProt-Swissprot human database using the in-built Andromeda data-search engine. Trypsin was selected as the enzyme (up to 2 missed cleavages), carboamidomethylation (C) as fixed modification and Deamidated (NQ) and Oxidation (M) as variable modifications. Protein false discovery rate was set at 1%. Data were quantified using the label-free quantitation (LFQ) and the Intensity-Based Absolute Quantification (iBAQ) parameter was enabled. Match between runs was not selected. MaxQuant protein group output was further analysed using Perseus (1.6.2.2). For the TurboID-cyclin F dataset (n of 3 per condition), a log2 transformation was applied to intensities prior to normalision by median subtraction. A 20% total valid number filter was then applied. Missing values were imputated (following the normal distribution). A two-sample student t-test was applied in combination with a Permutation-FDR correction (5%). For the EXO1 WT vs T824A interactome dataset (n of 1 per condition), a log2 transformation was used on iBAQ values and missing values were replaced by a constant equal to 5.

The raw MS data included in this paper will be deposited to the Proteome eXchange Consortium via the PRIDE partner repository with the dataset identifier^48^.

### CRISPR Knockout

Stable knockout was generated by co-transfecting Cas9 protein and 3 sgRNAs targeting the same gene, designed by Synthego, using Lipofectamine CRISPR Max (Life Technologies, CMAX00001) according to the protocol here: https://www.synthego.com/products/crispr-kits/synthetic-sgrna. 4 days after transfection, cells were trypsinised and seeded as single cells in 96-well plates to isolate single clones. Proliferative populations were then trypsinised and expanded in bigger vessels until there were sufficient cells for immunoblotting. Clones that showed clear knockout were further validated by gDNA extraction and PCR amplification of the exon targeted by the sgRNAs. The PCR product was ligated into pCR™4 vector using TOPO™ TA Cloning™ Kit (Invitrogen, 450030). After transforming into DH5α competent cells, 10 colonies were picked and sent for Sanger sequencing. In LN229 populations, cells were co-transfected with Cas9 protein and Synthego sgRNAs and directly subjected to treatment and western blotting 4 days after transfection. Although single clones were not isolated, knockout efficiency in the mixed population proved adequate (Figure 3D).

### Transfection, Immunoprecipitation, and Western Blotting

1.5 million HEK293T cells were seeded into each of the 10 cm petri dishes 24 hours before plasmid transfection. For each 10 cm dish, the PEI transfection was performed by vigorous vortexing of 5 μg plasmid DNA with 15 μl 2.5 mg/ml PEI in 400 μl DMEM (without FBS or antibiotics) for 15 seconds to mix, incubationfor 15 minutes at room temperature, then addition to cells cultured in 10 ml complete media in a drop-wise fashion. PEI (linear polyethylenimine) was purchased from Polysciences Inc (23966). 2.5 mg/ml PEI stock was made in 20 mM HEPES, 150 mM NaCl, pH 7.4, and filtered. 24 hours after transfection, cells were washed twice with PBS and harvested. The cell pellets were stored at −80 ℃ or lysed directly for experiments.

Cell pellets harvested for immunoprecipitation (IP) or western blotting (WB) not aiming to detect DNA damage markers were lysed in lysis buffer containing 50 mM Tris-HCl pH 7.5, 150 mM NaCl, 1 mM EDTA, 5 mM MgCl_2_, and 0.1% Nonidet P-40, supplemented with protease inhibitor cocktail (Sigma-Aldrich, P8340), 200 μM PMSF (Santa Cruz, sc-482875), and two phosphatase inhibitors – 20 mM beta-glycerophosphate (Sigma-Aldrich, G5422) and 1 µM okadaic acid (Cayman Chemical, 10011490-50 ug-CAY). After lysing on ice for 10 minutes, the insoluble fraction (mostly DNA and DNA-bound proteins) was removed via centrifugation at 20,000 g at 4 ℃ for 15 min, and the supernatant was carefully transferred to new Eppendorf tubes without disturbing the insoluble fraction. The protein concentration of the supernatant was measured using the modified Lowry assay (DC^TM^ Protein Assay Kit, BIO-RAD, 5000111). The same amount of total protein was mixed with Laemmli sample buffer (Invitrogen, NP0008) then used for each IP (0.5-1 mg per pulldown in general) or direct WB (10-20 μg per lane in general). For IP, 10 μl Flag M2 beads (Sigma-Aldrich, A2220-5ML) or 15 μl HA beads (Sigma-Aldrich, E6779-1ML) were washed 3 times with lysis buffer and added to the cell lysates to incubate for 3 hours on a roller at 4°C. After incubation, the beads were collected by centrifugation and washed with inhibitor-containing lysis buffer 5 times before being mixed with 20-40 μl 1× Laemmli sample buffer (diluted from Invitrogen, NP0008) supplemented with β-mercaptoethanol, and boiled for 10 minutes at 95°C. The boiled supernatant can then be used for WB analysis.

For experiments detecting DNA damage markers (which could be tightly chromatin-bound), cell pellets were harvested, homogenized, boiled directly in 2% SDS buffer (350 mM Bis-Tris pH 6.8, 20% glycerol, and 2% SDS), and then sonicated. Protein concentration was assessed using a BCA protein kit (Thermo Scientific, 23227). Cell lysate was prepared for WB as indicated above and resolved in 4-12% gradient Bis-Tris gels, transferred onto nitrocellulose membrane (Amersham, 10600006) and immunoblotted. WB results were visualized on X-ray film or using the iBright™ FL1500 Imaging System (Invitrogen, A44241).

### Cell Cycle Synchronisation via Double Thymidine Release (DTR)

500,000 cells were seeded into each 10 cm dish. For HeLa parental and HeLa *CCNF* K/O cell lines, cells were seeded directly into the first thymidine block (2 mM final concentration) and cultured for 16 h. After being washed 3 times with PBS and once with complete media, cells were released into fresh complete media for 8 h. A second thymidine block was then performed with the same conditions as the first. 16 hours later, 0 h time point samples were collected before the rest of the cells were washed and released into fresh complete media. Cells were collected at different time points as indicated. At the 7 h time point, 200 nM nocodazole was added to the uncollected cells to prevent cells from entering the next cell cycle. For cell cycle synchronisation in LN229 cells, all conditions were kept the same except the duration of the thymidine blocks, which changed from 16 h to 24 h.

### Cycloheximide (CHX) Chase

250,000 cells were seeded into each well of a 6-well plate. 16 hours later, cells were cultured with 50 μg/ml cycloheximide for various durations as indicated in the figures, to block ribosomal protein synthesis for assessment of protein stability. Cells were collected by scraping and washed with PBS twice before being subjected immediately to WB for protein half-life estimation.

### *In vivo* ubiquitination Assay

2 million HEK293T cells were seeded into each 10 cm dish 24 h before being transfected with plasmids as indicated in each experiment (typically, for each 10 cm dish, 1 μg substrate overexpression plasmid, 2 μg E3 overexpression plasmid, or 3 μg ubiquitin overexpression plasmid were co-transfected). Cells were cultured for another 24 hours before being harvested. 4 hours before the harvest, MG132 was added to a final concentration of 10 μM to block proteasomal degradation so that ubiquitination events were enriched. Cells were collected by scrapping and washed with PBS once, then thoroughly lysed and boiled in 300 μl ubiquitin lysis buffer (2% SDS, 150 mM NaCl, 10 mM Tris–HCl pH 7.4). After cooling to room temperature, cell lysates were subjected to sonication until they lost their viscosity. Lysates were then boiled again and centrifuged at 17,000 x g for 10 min. 20 μl supernatant was preserved as input for each sample. The rest of the supernatant was diluted 20 times with dilution buffer (10 mM Tris–HCl pH 7.4, 150 mM NaCl, 2 mM EDTA, 1% Triton x-100) and processed by immunoprecipitation of the substrate. After 16 h on a roller at 4 ℃, beads were collected via centrifugation at 2000 rpm for 1 minute and washed 5 times with 1 ml wash buffer (10 mM Tris–HCl pH 7.4, 1 M NaCl, 1 mM EDTA, 1% Nonidet P-40) before being mixed with 50 μl Laemmli sample buffer, boiled and subjected to WB.

### Ubiquitin Binding Entities (UBE) Pulldown Assay

Prior to harvesting the cells, GST-UBA (glutathione S-transferase) was first conjugated to glutathione sepharose beads (Cytiva, GE17-0756-01) in UBE lysis buffer (19 mM NaH_2_PO_4_, 81 mM Na_2_HPO_4_, pH 7.4, 1% Nonidet P-40, 2 mM EDTA, supplemented with protease inhibitor cocktail, PMSF, and phosphatase inhibitors as mentioned in the immunoprecipitation section, 50 mM N-Ethylmaleimide (NEM, Sigma-Aldrich, E3876-5G), 5 mM 1,10-Phenanthroline (Scientific Laboratory Supplies, CHE2730), and 50 μM PR-619 (ApexBio, A821) for at least 4 hours at 4 ℃ on a roller. For each pulldown, 100 μg recombinant GST-UBA was conjugated to 20 μl of washed glutathione beads. Cells were treated with 10 μM MG132 for 4 hours before being harvested by scrapping and centrifugation. Cell pellets were washed twice with PBS, then directly lysed in freshly prepared UBE lysis buffer. After incubation on ice for 10 minutes, lysates were subjected to centrifugation at 14,000 rpm at 4 ℃ for 15 minutes. After the protein concentration was measured via Lowry assay using DC^TM^ Protein Assay Kit (BIO-RAD, 5000111), the same amount of total protein was used for each pulldown (2-3 mg per pulldown). GST-UBA-conjugated beads were added to the lysate and incubated on a roller at 4 ℃ overnight, then collected, washed using UBE lysis buffer, mixed with 1× Laemmli sample buffer and boiled in the same way as described in the immunoprecipitation section. The supernatant was then used for WB. Besides the protein of interest, total ubiquitin was probed as a loading control for UBE pulldown experiments.

### *In vitro* Dephosphorylation

*In vitro* dephosphorylation was performed using lambda phosphatase kit (New England Biolabs, P0753S). Following the immunoprecipitation step, beads were washed 3 times with lysis buffer containing only protease inhibitors but not phosphatase inhibitors, and once with the 1 X lambda phosphatase buffer provided in the kit. The wash buffer was removed, and the beads were then resuspended in 100 μl 1 X lambda phosphatase buffer containing 1 mM MnCl_2_ and 400 U lambda phosphatase and incubated at 30 ℃, 800 rpm, for 1 h on an orbital shaker to remove phosphorylation on all proteins bound to the beads. After the incubation, the beads were washed 3 times with 1 X lambda phosphatase buffer before being used for WB or for *in vitro* phosphorylation assay.

### *In vitro* Phosphorylation Assay

For *in vitro* kinase assays, EGFP-tagged EXO1 was overexpressed in HEK293T cells and purified by immunoprecipitation using GFP-Trap beads (Proteintech, gta-20). Before being used as the substrate, EGFP-EXO1-bound beads were first dephosphorylated by lambda phosphatase (New England BioLabs, P0753S) to remove all existing phosphorylation. After washing off the lambda phosphatase extensively with kinase buffer (Cell Signaling Technologies, 9802S), beads were incubated for 30 min at 30 ℃ in 100 ng CDK1/2-Cyclin A/B/E in kinase buffer supplemented with 200 μM ATP. The reactions were stopped by with the addition of 4× Laemmli sample buffer and incubation at 95 ℃ for 5 minutes. EXO1 T824 phosphorylation was visualized by SDS-PAGE and WB using the custom-made site-specific phosphorylation antibody described below.

### Generation of Site-Specific Phosphorylation Antibody Targeting pT824 EXO1

Antibodies were generated by YenZym Antibodies LLC. The following pThr824–EXO1 peptide was synthesized and conjugated to a carrier protein through the N-terminal cysteine residue (Ahx is added as a linker). As a negative control for validation experiments, peptide with the same sequence but no phosphorylation was also synthesized. Two rabbits were used for immunisation. 21 days after immunisation, their serum was harvested and subjected to affinity absorption and ELISA validation.

pThr824 EXO1 peptide:

C-Ahx-RDNIQLp<u>T</u>PEAEED-amide (amino acid residues from 818 to 830)

Non-phosphorylated Thr824 EXO1 peptide:

C-Ahx-RDNIQL<u>T</u>PEAEED-amide

### *In situ* Detection of ssDNA

The protocol used in this study was a modification of the one described by Caron et al.^49^. In short, to visualize and quantify ssDNA generated by DNA resection, EXO1 expression in LN229 cells containing pCW57.1-EXO1 WT or R842A was induced for 3 days in complete media containing 1 μg/ml doxycycline before 300,000 cells per well were seeded in a 6-well plate pre-filled with sterile glass coverslips (also with doxycycline). After the cells attached to the coverslips, 10 μM BrdU (Cayman Chemical, CAY15580-500 mg) was added and the cells were incubated for 24 hours. BrdU-containing media was then replaced with fresh complete media containing no BrdU, but with doxycycline still present. Cells were then treated with 10 Gy IR and allowed to recover for 3 hours before being subjected to *in situ* fractionation on ice as follows: 10 minutes in pre-extraction buffer 1 (10 mM PIPES, pH 7.0, 300 mM sucrose, 100 mM NaCl, 3 mM MgCl2, and 0.5% Triton-X100), then 10 minutes in pre-extraction buffer 2 (10 mM Tris-HCl pH 7.5, 10 mM NaCl, 3 mM MgCl2, 1% Nonidet P-40, and 0.5% sodium deoxycholate). Cells on coverslips were washed 3 times with PBS before being fixed with 4% paraformaldehyde for 15 min at room temperature. The cells were then washed with PBS, and blocked and permeabilized in 3% BSA and 0.5% Triton X-100 dissolved in PBS for 30 min. They were incubated overnight at 4 °C with mouse anti-BrdU antibody (1:500 dilution, BD Biosciences, 347580) under non-denaturing conditions followed by anti-mouse secondary antibody (1:1000 dilution, Invitrogen, A-11004) for 1 hour. As the anti-BrdU antibody detects BrdU incorporated into ssDNA but not dsDNA, the BrdU signal detected via this protocol is a good reflection of ssDNA generated by DNA resection. Following the antibody incubation, the cells were washed 5 times in 0.2% Triton X-100 in PBS before being mounted onto slides using VECTASHIELD antifade mounting medium (H-1000) supplemented with 5 μg/ml DAPI and visualized using a Zeiss LSM 780 Confocal Microscope.

### SMART Assay

On average, ∼1,000,000 cells per 10 cm dish were seeded for each sample in the experiment. After the cells attached, 10 μM BrdU (Cayman Chemical, CAY15580-500 mg) was added to the cell culture and incubated for 24 hours. BrdU-containing media was then switched to fresh complete media plus doxycycline but not BrdU. Cells were treated with 10 Gy IR and allowed to recover for 3 hours before being harvested by trypsinisation, counted, and embedded in low-melting-point agarose gel plugs. About 250,000 cells were used for each plug. DNA extraction in the plug, agarose digestion of the plug, and combing were performed following the instructions in the kit manual of FiberPrep DNA Extraction Kit (Genomic Vision, EXT-001A). BrdU was detected by 1-hour incubation of mouse anti-BrdU antibody (1:10 dilution, BD Biosciences, 347580) at 37 °C followed by incubation for 1 h in the Cy3.5-conjugated anti-mouse secondary antibody (1:10 dilution, Abcam, AB6946) at 37 °C under non-denaturing conditions. Coverslips were scanned using the automated FibreVision®S scanner. Results were quantified using FiberStudio.

### Anaphase UFB Detection and Micronuclei Counting

After treating cells with doxycycline for 3 days, cells were seeded at 200,000 cells per well in 6-well plates filled with sterile coverslips. 24 hours after seeding, cells were treated with 10 Gy IR and allowed to recover for 12 hours. Cells on coverslips were fixed and permeabilised with UFB buffer (4% PFA in 20 mM PIPES at pH 6.8, 1 mM MgCl2, 10 mM EGTA, and 0.2% Triton X-100) for 10 minutes at room temperature. Samples were then washed with PBS containing 0.2% Triton for 5 minutes and blocked by 3% BSA for 30 minutes. Primary antibodies (rabbit anti-BLM from Abcam (ab2179) and mouse anti-RPA32 from Abcam (ab2175)) diluted in 3% BSA were then applied on top of each coverslip in a wet chamber and incubated at 4 °C overnight. Next day, the unbound primary antibodies were washed off 3 times using 0.2% Triton in PBS before the coverslips were incubated with secondary antibodies (A-11004 and A-11008 from Invitrogen) diluted in 3% BSA for 2 hours at room temperature. Coverslips were washed 3 times with PBS before being mounted onto glass slides using VECTASHIELD antifade mounting medium (H-1000) supplemented with 5 μg/ml DAPI. UFBs and micronuclei were visualized and counted using a Zeiss LSM 780 Confocal Microscope.

### Mitotic spreading

EXO1 expression in hTERT RPE-1 cells containing pCW57.1-EXO1 WT or R842A constructs was induced for 3 days in complete media containing 1 μg/ml doxycycline before 500,000 cells per 10 cm dish were seeded for the experiment (also with doxycycline). 24 hours after reseeding, cells were treated with 10 Gy IR and allowed to recover for 3 days. 3 hours before harvesting, cells were also treated with 0.02 μg/ml colcemid (GIBCO, 15212012) to enrich metaphase events. At the end of the colcemid treatment, media was collected to preserve the floating mitotic cells, and the adherent cells were harvested via trypsinisation, resuspended thoroughly in 10 ml hypotonic solution (75 mM KCl in water), and incubated at 37 ℃ for 30 minutes. All cells were then collected via centrifugation at 200 xg for 10 minutes. The supernatant was removed until ∼300 μl was left. The cells were then thoroughly resuspended in the remaining supernatant by flicking before being incubated on ice for 30 minutes in 5 ml freshly-made fixative (methanol: glacial acetic acid at 3:1 ratio), added in a drop-wise fashion,. The now fixed cells were once again centrifuged then resuspended in 300 μl supernatant followed by 5 ml fixative to substitute all hypotonic buffer with fixative. After repeating the substitution once more, cells were thoroughly resuspended in 200 μl fixative and kept on ice before 30-50 μl of the cell suspension was dropped on a clean glass slide from a 10-15 cm vertical distance for metaphase spreading. The slide was then airdried for 2 hours before the application of VECTASHIELD antifade mounting medium (H-1000) supplemented with 5 μg/ml DAPI and subsequent visualization using a Zeiss LSM 780 Confocal Microscope.

### Cyclin F-EXO1 model

A computational protein-protein docking method was used to predict the atomic interactions between cyclin F and EXO1. ‘COSMIC^2^’ was used for structure prediction. The inputs included Accession number NP_001752.2 for cyclin F and the C terminal peptide sequence ‘LEPEAEEDIFNKPECGRVQRAIFQ’ for EXO1. The parameters were set to consider ‘full database’, ‘multimer’, and ‘no relaxed’ models. Five multimer models were generated and confidence scores of iptm+ipm were compared. ChimeraX was used to visualize the chosen model. The model had an iptm+ptm value of 0.7984765224933627.

## ACKNOWLEDGMENTS

This study was made possible thanks to the support of a Medical Research Council (MRC) grant MC_UU_00001/7 to V.D’A. We acknowledge further support by a John Fell (133/075) and Wellcome Trust grant (097813/Z/11/Z) to B.M.K. and F.M.B, and we acknowledge support from European Research Council Award microC 772970. The TurboID proximity labelling expression vector is developed by Dr. Kilian Huber’s lab. We are grateful to receive it as a kind gift. We thank Dr. Jagat Chauhan for providing help in data trimming and cleaning in prior to our screen analysis. We also would like to thank Dr. Zhuoyao Chen and Dr. Alex Bullock for kindly sharing GST-UBA protein for the UBA assay. Lastly, we would like to thank Dr. Varun Gopala Krishna and Katherine Ferris for proof-reading the paper.

## AUTHOR CONTRIBUTIONS

Conceptualization and experiment design, H.Y. and V.DA.; CRISPR screen and major methodology, H.Y.; Phosphorylation study, S.F.; TurboID cell line generation and Mass spectrometry sample preparation P.S.; molecular docking, E.Y.B.; immunofluorescence data acquisition and analysis, H.Y. and Y.J.; Bioinformatic analysis, F.B. and A.V.N.; next generation sequencing, D.B.; Mass spectrometry, R.F., I.V., and B.K.; writing and original draft, H.Y., V.DA.; proofreading & editing, all authors; supervision, V.DA..

## INCLUSION AND DIVERSITY

We support inclusive, diverse, and equitable conduct of research. We worked to ensure ethnic or other types of diversity in the recruitment of human subjects. We worked to ensure sex balance in the selection of non-human subjects. One or more of the authors of this paper self-identifies as an under-represented ethnic minority in their field of research or within their geographical location. One or more of the authors of this paper self identifies as a gender minority in their field of research. One or more of the authors of this paper self-identifies as a member of the LGBTQIA+ community. One or more of the authors of this paper received support from a program designed to increase minority representation in their field of research. We also avoided ‘‘helicopter science’’ practices by including the participating local contributors from the region where we conducted the research as authors on the paper.

---

**Figure S1.**
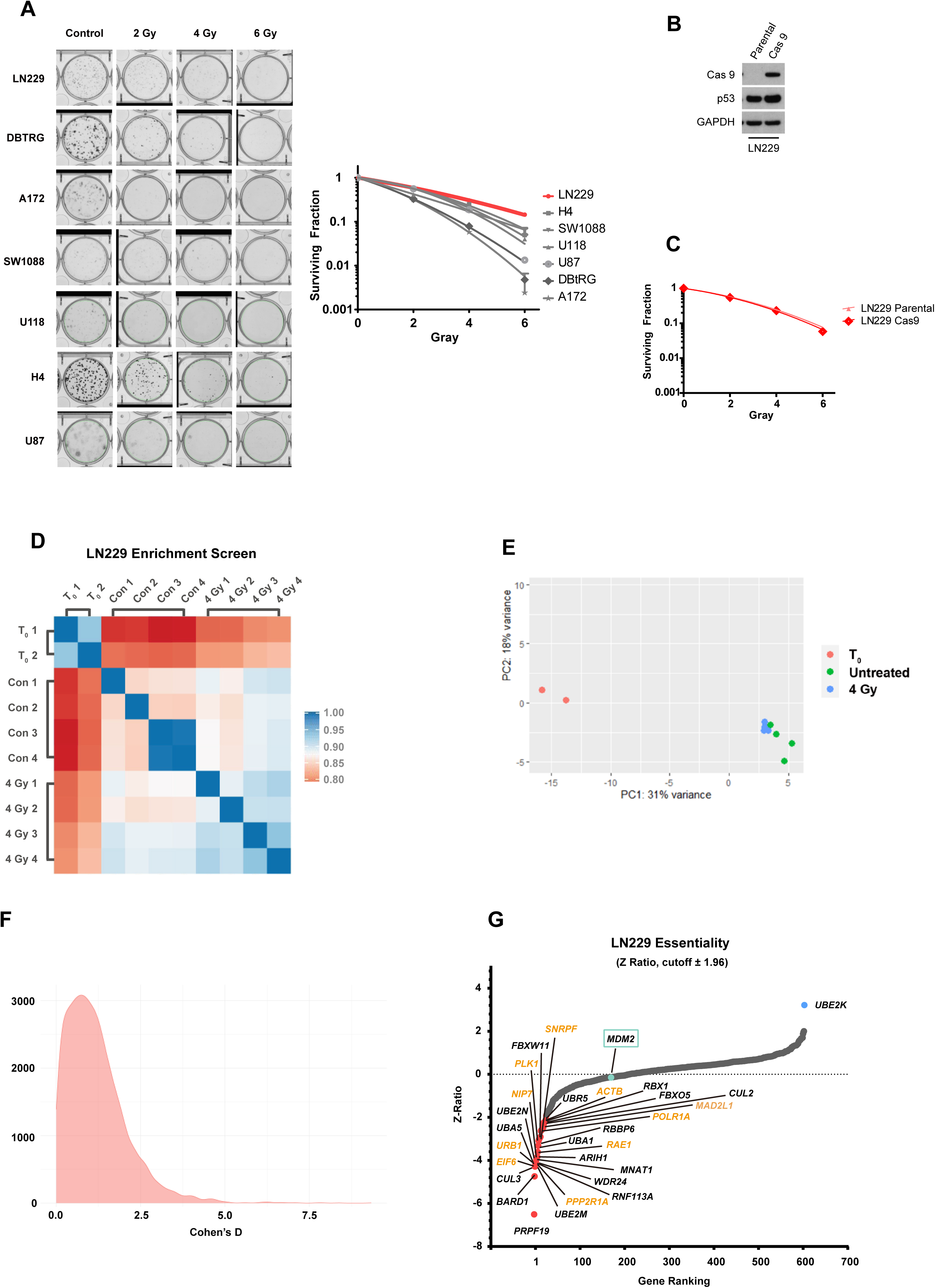
Optimisation and Quality Control of the CRISPR Screen. A. The 7 GBM cell lines as indicated were seeded for colony formation assay and challenged with the indicated dose of IR. 14 days after IR, cells were stained with crystal violet and counted. Error bars represent standard deviations of three biological replicates. B. Immunoblotting of LN229 single clone derived cells expressing Cas9 after lentiviral transduction; C. Cells in B. were seeded for colony formation assay and challenged with the indicated dose of IR. 14 days after IR, cells were stained with crystal violet and counted. Error bars represent standard deviations of three biological replicates. D. Rank correlation of normalized sgRNA read counts between biological replicates and treatment conditions of the screen; E. Principal component analysis (PCA) of different samples of the screen. Sample T0 is in orange, quadruplicated samples for untreated condition are in green, and 4 Gray IR treated samples are in blue; F. Frequency plot depicting the frequency effect size Cohen’s d values between treated and untreated samples for each CRISPR guide. G. Ranked genes using Z Ratio after comparing untreated LN229 to T0. Genes with negative Z ratio are considered essential genes (in red). Gene with positive Z ratio are in blue. ±1.96 Z-ratio value was used as cut-offs. Gene names in orange indicates core essential positive control genes provided in the sgRNA library.

**Figure S2.**
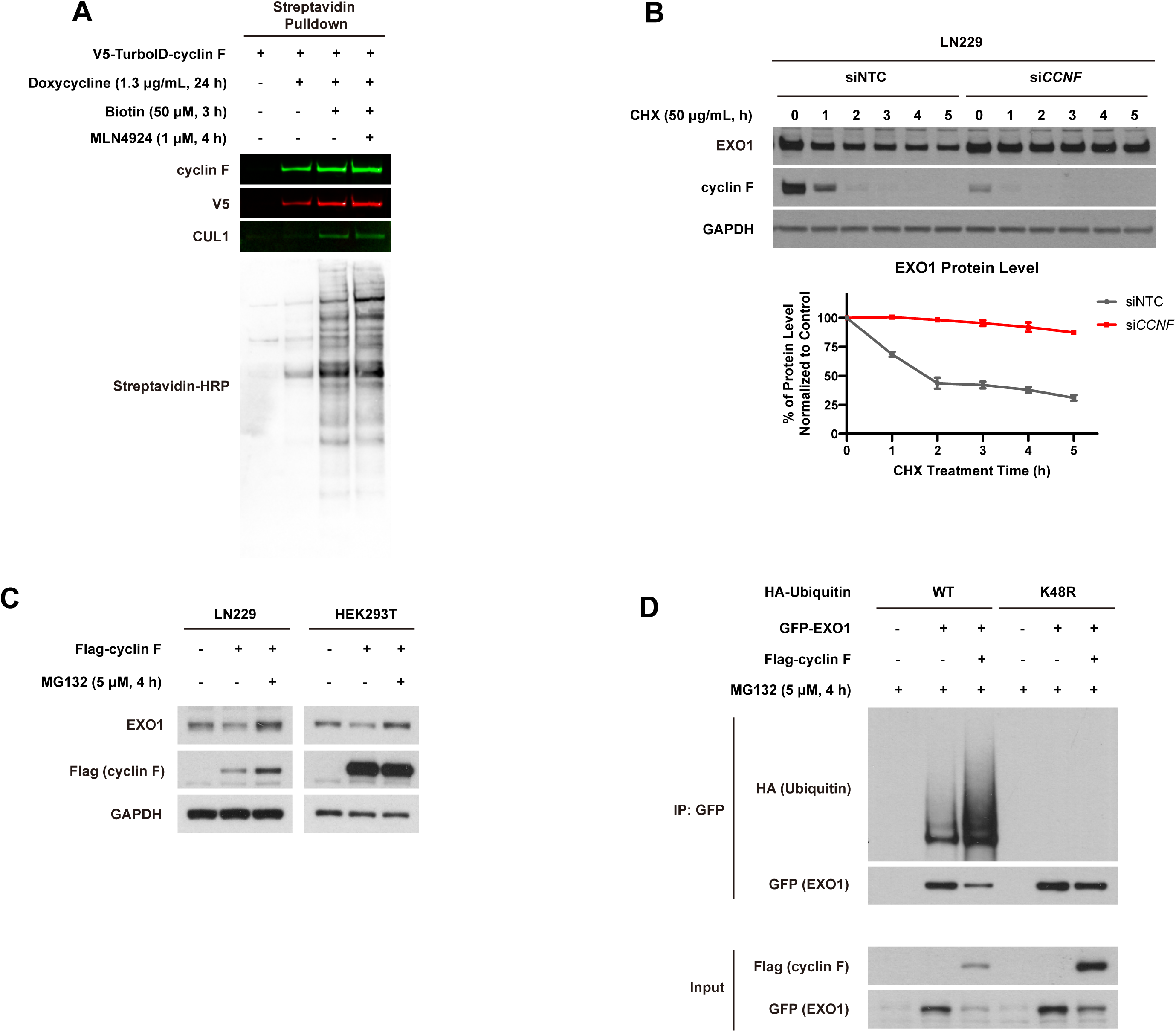
SCF^cyclin^ ^F^ Ubiquitinates and Degrades EXO1. A. Immunoblotting after isolation of biotinylated proteins from HEK293T cells expressing TurboID-cyclin F after induction with doxycycline for 24 h. Cells were doxycycline induced and treated with biotin and MLN4924 as indicated. Biotinylated proteins were detected by HRP-conjugated streptavidin- *bottom panel*. B. Immunoblotting after transfection of LN229 cells with non -targeting control (NTC) siRNA or siRNA targeting *CCNF* treated with Cycloheximide (CHX) for the indicated time -*upper panel*. Relative quantification of EXO1 protein levels in cells after normalisation with EXO1 levels at T0 for each cell line - *bottom panel*. C. Immunoblotting of LN229 and HEK293T after transient transfection of GFP-cyclin F and treatment with MG132 as indicated. D. Immunoblotting after expression of GFP-EXO1, Flag-cyclin F, HA-ubiquitin wild-type or K48R mutant in HEK293T. GFP-EXO1 is isolated *via* GFP beads pulldown before immunoblotting. Input samples before immunoprecipitation are indicated.

**Figure S3.**
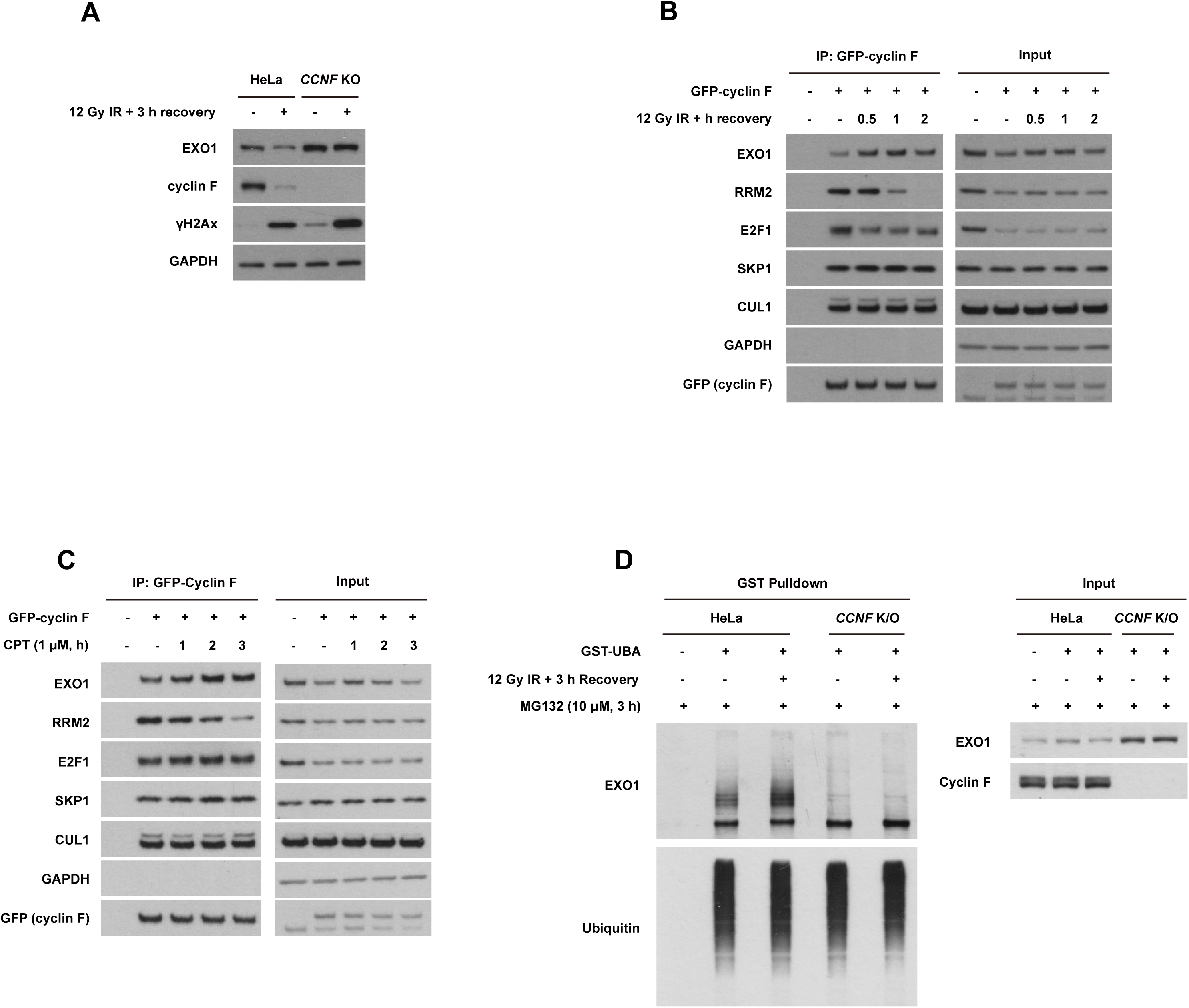
SCF^cyclin^ ^F^-Mediated EXO1 ubiquitination is induced by DNA damage. A. Immunoblotting of HeLa and *CCNF* K/O after treatment with IR and MG132. B. Immunoblotting after immunoprecipitation of GFP-cyclin F in HEK293T cells treated with IR. *left panel*. Input samples before immunoprecipitation are indicated – *right panel*. C. Immunoblotting after immunoprecipitation of GFP-cyclin F in HEK293T cells treated with or without Camptothecin (CPT) for indicated durations. D. Immunoblotting after isolation of endogenous ubiquitinated proteins using recombinant GST-tagged UBA domain of UBIQLN protein in HeLa or *CCNF* K/O treated with IR and MG132 as indicated.

**Figure S4.**
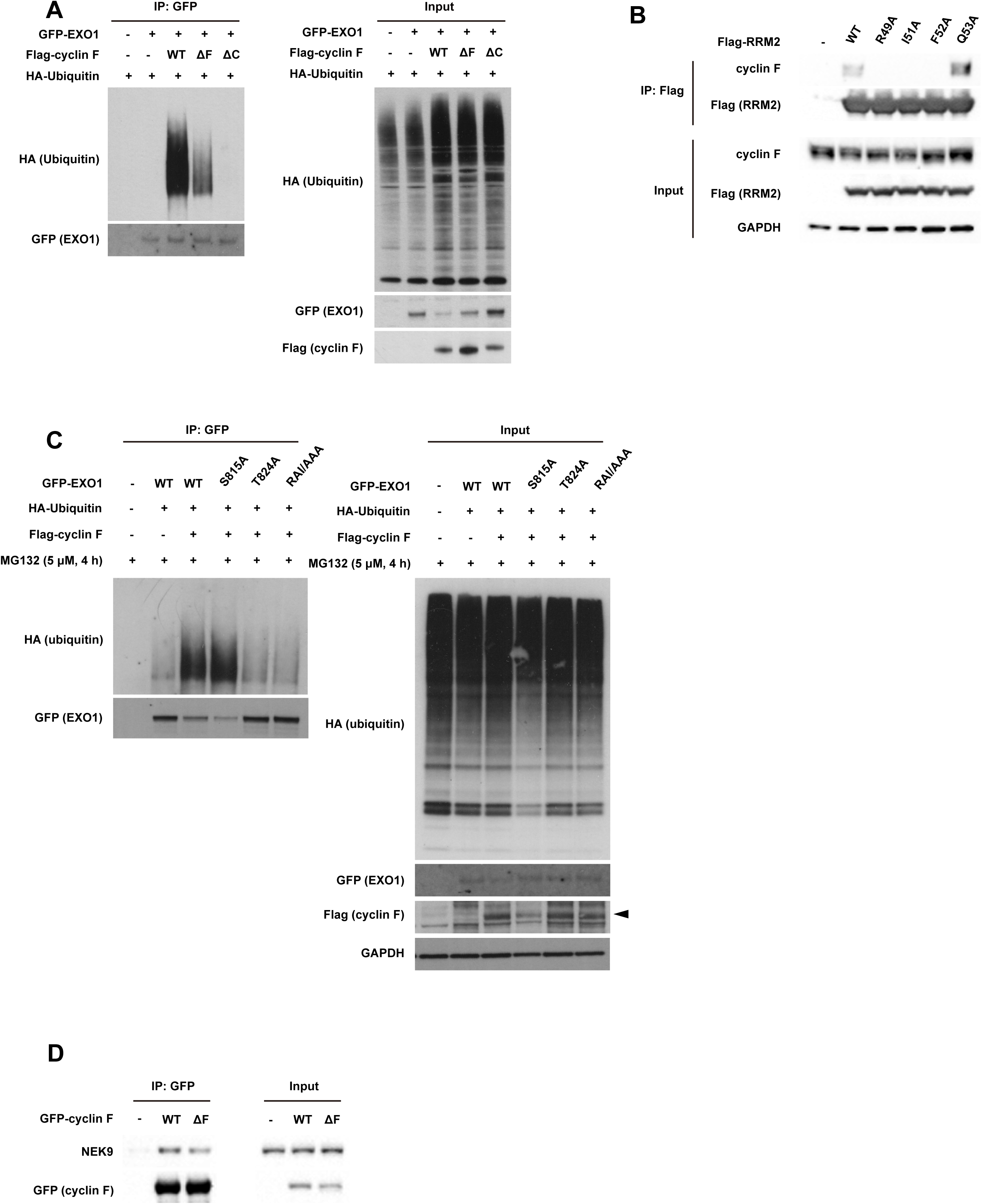
Identification of Domain and Motif for Cyclin F-EXO1 Interaction. A. Immunoblotting after expression of Flag-cyclin F WT, Flag-cyclin F ΔF or Flag-cyclin F ΔC and Flag-EXO1 with HA-ubiquitin in HEK293T. GFP-EXO1 is isolated *via* GFP beads pulldown before immunoblotting. B. Immunoblotting of immunoprecipitated Flag-tagged RRM2 WT, R49A, I51A, F52A and Q53A mutant from HEK293T. C. Immunoblotting after expression of GFP-EXO1 WT, GFP-EXO1 S185A, GFP-EXO1 T824A or GFP-EXO1 RAI842AAA and Flag-cyclin F WT, with HA-ubiquitin in HEK293T. GFP-EXO1 is isolated *via* GFP beads pulldown before immunoblotting. D. Immunoblotting of immunoprecipitated GFP-tagged cyclin F or GFP-cyclin F ΔF in HEK293T.

**Figure S5.**
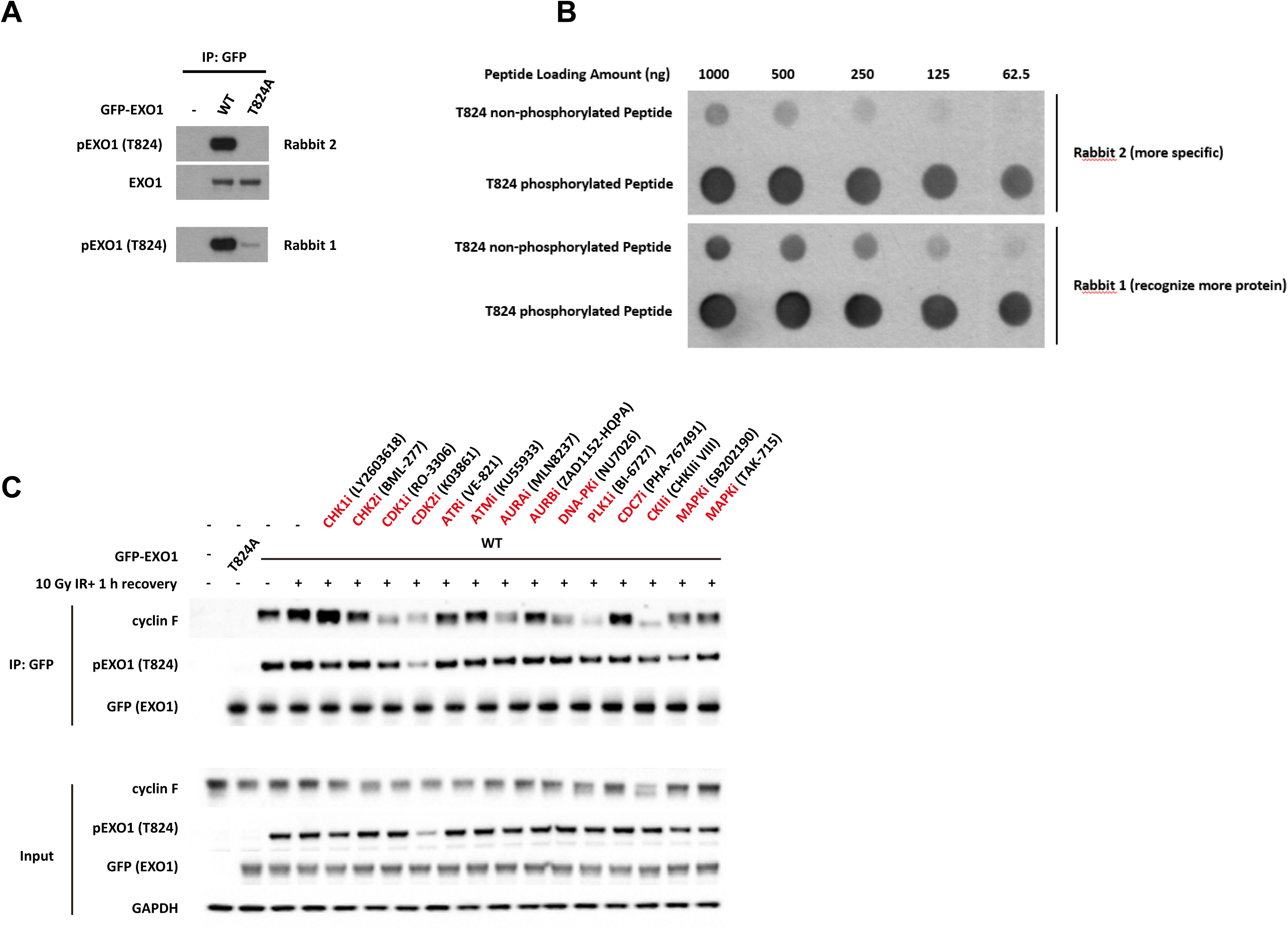
T824 Phosphorylation is Crucial for SCF^cyclin^ ^F^-Mediated EXO1 regulation. A. Immunoblotting of immunoprecipitated GFP-EXO1 WT or GFP-EXO1 T824A from HEK293T. B. Different amount of T824 phosphorylated or non-phosphorylated EXO1 peptides (indicate sequences) were spotted on nitrocellulose membrane. After air-drying, the membrane is immunoblotted for pT824 EXO1 using site-specific antibodies generated from immunising two rabbits (1,2 as indicated); C. Immunoblot of immunoprecipitated GFP-EXO1 from HEK293T cells after treatment with the indicate kinase inhibitors.

**Figure S6.**
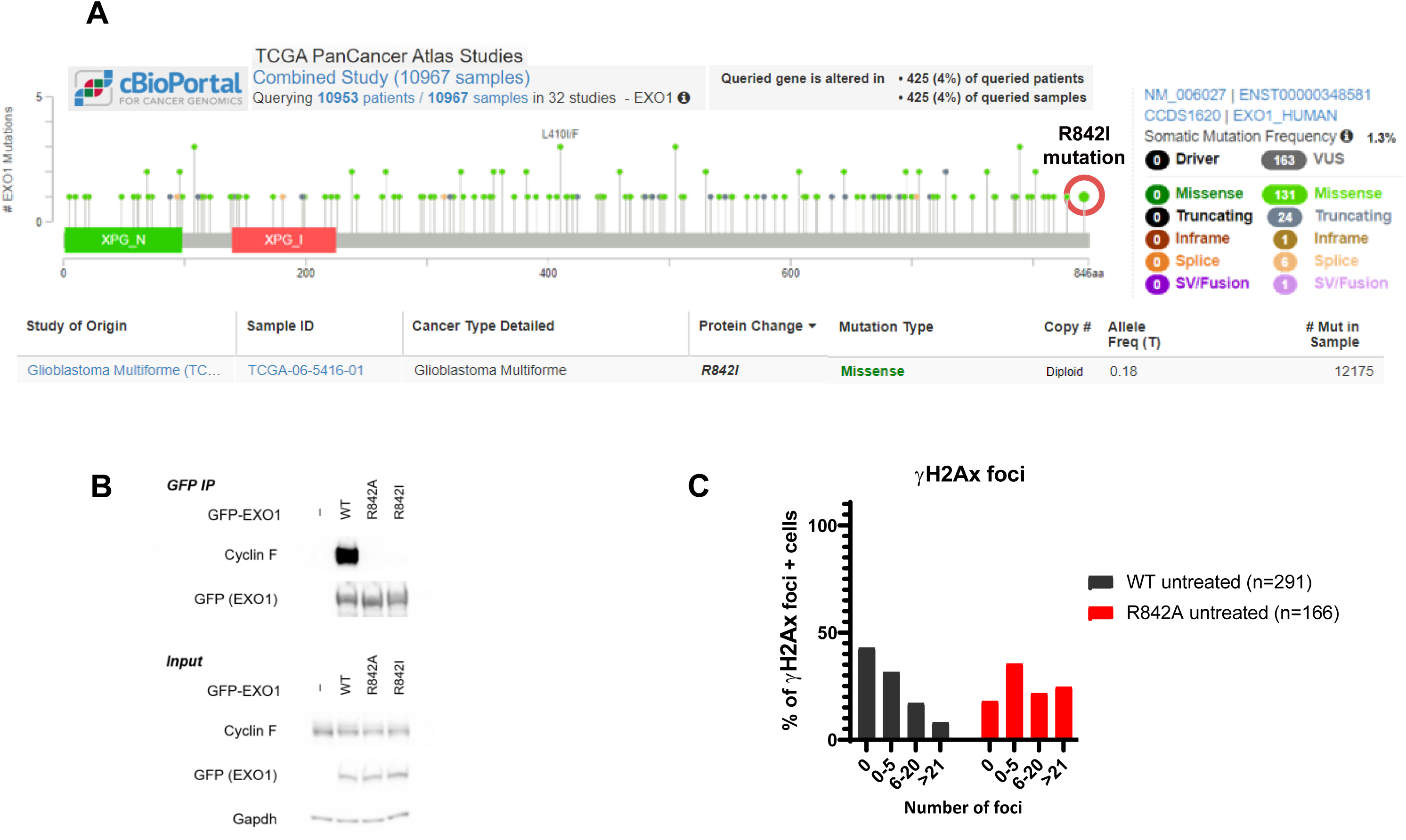
R842 Mutation is Found in one GBM Patient. A. Snapshot of EXO1 mutations identified in the cBioportal encompassing 10932 samples in 32 studies. The mutation identified in one case of glioblastoma multiforme within the RxIF motif is highlighted. B. Immunoblotting of immunoprecipitated GFP-EXO1 WT, GFP-EXO1 R842A, GFP-EXO1 R842I. C. Quantification of gH2Ax foci in cells expressing HA-EXO1 WT or HA-EXO1 R842A after IR treatment at 10 Gy at the indicated time.

**Figure S7.**
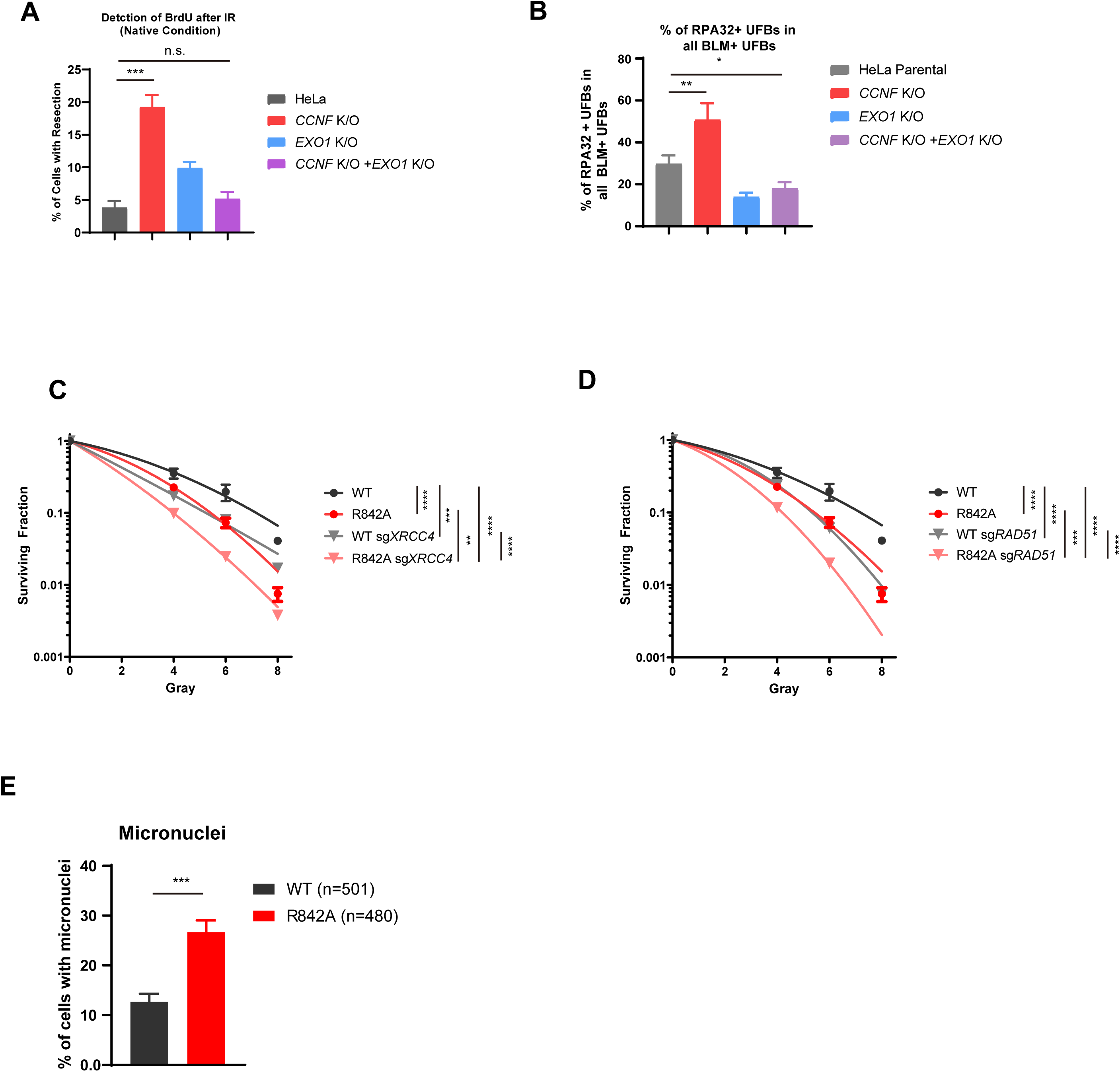
*CCNF* K/O or EXO1 R842A Mutation Both Lead to Genome Instability. A. Quantification of BrdU signal in HeLa parental cells, HeLa *CCNF* K/O, HeLa *EXO1* K/O and HeLa *CCNF* K/O *EXO1* K/O. Cells were seeded for 50% density and labelled with 10 μM BrdU for 24 hours before being treated with 10 Gy IR and allowed to recover for 3 hours. Error bars represent standard deviations of three biological replicates. Statistical analysis was done using two-tailed unpaired t test. *** indicates P ⩽ 0.001, n.s. = not significant; B. Quantification of RPA32 positive anaphase ultra-fine bridges (UFBs) in HeLa parental cells, HeLa *CCNF* K/O, HeLa *EXO1* K/O and HeLa *CCNF* K/O *EXO1* K/O. Cells were treated with 10 Gy IR and allowed to recover for 12 hours before subjected to UFB detection. At least 50 events were quantified for each condition. Error bars represent standard deviation of three biological replicates. Statistical analysis was done using two-tailed unpaired t test. ** indicates P ⩽ 0.01, n.s. indicates not significant. C. LN229 expressing HA-EXO1 WT or HA-EXO1 R842A and transfected with a control sgRNA or sgRNA targeting *XRCC4* as indicated were seeded for colony formation assay and challenged with the indicated dose of IR. 14 days after IR, cells were stained with crystal violet and counted. Error bars represent standard deviations of three biological replicates. D. LN229 expressing HA-EXO1 WT or HA-EXO1 R842A and transfected with a control sgRNA or sgRNA targeting *RAD51* as indicated were seeded for colony formation assay and challenged with the indicated dose of IR. 14 days after IR, cells were stained with crystal violet and counted. Error bars represent standard deviations of three biological replicates. E. Quantification of micronuclei in LN229 expressing HA-EXO1 WT or HA-EXO1 R842A. Error bars indicate standard deviations of three biological replicates. n represents total events quantified in all three replicates. Statistical analysis was done using two-tailed unpaired t test. *** indicates P ⩽ 0.001.

## References

1. Jackson, S.P., and Bartek, J. (2009). The DNA-damage response in human biology and disease. Nature 461, 1071–1078. 10.1038/nature08467.

2. Schwertman, P., Bekker-Jensen, S., and Mailand, N. (2016). Regulation of DNA double-strand break repair by ubiquitin and ubiquitin-like modifiers. Nat Rev Mol Cell Biol 17, 379–394. 10.1038/nrm.2016.58.

3. Chapman, J.R., Taylor, M.R., and Boulton, S.J. (2012). Playing the end game: DNA double-strand break repair pathway choice. Mol Cell 47, 497–510. 10.1016/j.molcel.2012.07.029.

4. Bennardo, N., Cheng, A., Huang, N., and Stark, J.M. (2008). Alternative-NHEJ is a mechanistically distinct pathway of mammalian chromosome break repair. PLoS Genet 4, e1000110. 10.1371/journal.pgen.1000110.

5. Bhargava, R., Onyango, D.O., and Stark, J.M. (2016). Regulation of Single-Strand Annealing and its Role in Genome Maintenance. Trends Genet 32, 566–575. 10.1016/j.tig.2016.06.007.

6. Cejka, P., and Symington, L.S. (2021). DNA End Resection: Mechanism and Control. Annu Rev Genet 55, 285–307. 10.1146/annurev-genet-071719-020312.

7. Zhou, Y., Caron, P., Legube, G., and Paull, T.T. (2014). Quantitation of DNA double-strand break resection intermediates in human cells. Nucleic Acids Res 42, e19. 10.1093/nar/gkt1309.

8. Zierhut, C., and Diffley, J.F. (2008). Break dosage, cell cycle stage and DNA replication influence DNA double strand break response. EMBO J 27, 1875–1885. 10.1038/emboj.2008.111.

9. Tomimatsu, N., Mukherjee, B., Harris, J.L., Boffo, F.L., Hardebeck, M.C., Potts, P.R., Khanna, K.K., and Burma, S. (2017). DNA-damage-induced degradation of EXO1 exonuclease limits DNA end resection to ensure accurate DNA repair. J Biol Chem 292, 10779–10790. 10.1074/jbc.M116.772475.

10. Adeyemi, R.O., Willis, N.A., Elia, A.E.H., Clairmont, C., Li, S., Wu, X., D’Andrea, A.D., Scully, R., and Elledge, S.J. (2021). The Protexin complex counters resection on stalled forks to promote homologous recombination and crosslink repair. Mol Cell 81, 4440–4456 e4447. 10.1016/j.molcel.2021.09.008.

11. Elia, A.E., Boardman, A.P., Wang, D.C., Huttlin, E.L., Everley, R.A., Dephoure, N., Zhou, C., Koren, I., Gygi, S.P., and Elledge, S.J. (2015). Quantitative Proteomic Atlas of Ubiquitination and Acetylation in the DNA Damage Response. Mol Cell 59, 867–881. 10.1016/j.molcel.2015.05.006.

12. Sertic, S., Quadri, R., Lazzaro, F., and Muzi-Falconi, M. (2020). EXO1: A tightly regulated nuclease. DNA Repair (Amst) 93, 102929. 10.1016/j.dnarep.2020.102929.

13. Jackson, S.P., and Durocher, D. (2013). Regulation of DNA damage responses by ubiquitin and SUMO. Mol Cell 49, 795–807. 10.1016/j.molcel.2013.01.017.

14. Hustedt, N., and Durocher, D. (2016). The control of DNA repair by the cell cycle. Nat Cell Biol 19, 1–9. 10.1038/ncb3452.

15. D’Angiolella, V., Donato, V., Forrester, F.M., Jeong, Y.T., Pellacani, C., Kudo, Y., Saraf, A., Florens, L., Washburn, M.P., and Pagano, M. (2012). Cyclin F-mediated degradation of ribonucleotide reductase M2 controls genome integrity and DNA repair. Cell 149, 1023–1034. 10.1016/j.cell.2012.03.043.

16. D’Angiolella, V., Donato, V., Vijayakumar, S., Saraf, A., Florens, L., Washburn, M.P., Dynlacht, B., and Pagano, M. (2010). SCF(Cyclin F) controls centrosome homeostasis and mitotic fidelity through CP110 degradation. Nature 466, 138–142. 10.1038/nature09140.

17. Burdova, K., Yang, H., Faedda, R., Hume, S., Chauhan, J., Ebner, D., Kessler, B.M., Vendrell, I., Drewry, D.H., Wells, C.I., et al. (2019). E2F1 proteolysis via SCF-cyclin F underlies synthetic lethality between cyclin F loss and Chk1 inhibition. EMBO J 38, e101443. 10.15252/embj.2018101443.

18. Enrico, T.P., Stallaert, W., Wick, E.T., Ngoi, P., Wang, X., Rubin, S.M., Brown, N.G., Purvis, J.E., and Emanuele, M.J. (2021). Cyclin F drives proliferation through SCF-dependent degradation of the retinoblastoma-like tumor suppressor p130/RBL2. Elife 10. 10.7554/eLife.70691.

19. Cohen, J. (1977). Statistical power analysis for the behavioral sciences, Rev. Edition (Academic Press).

20. Stewart, G.S., Panier, S., Townsend, K., Al-Hakim, A.K., Kolas, N.K., Miller, E.S., Nakada, S., Ylanko, J., Olivarius, S., Mendez, M., et al. (2009). The RIDDLE syndrome protein mediates a ubiquitin-dependent signaling cascade at sites of DNA damage. Cell 136, 420–434. 10.1016/j.cell.2008.12.042.

21. Baranes-Bachar, K., Levy-Barda, A., Oehler, J., Reid, D.A., Soria-Bretones, I., Voss, T.C., Chung, D., Park, Y., Liu, C., Yoon, J.B., et al. (2018). The Ubiquitin E3/E4 Ligase UBE4A Adjusts Protein Ubiquitylation and Accumulation at Sites of DNA Damage, Facilitating Double-Strand Break Repair. Mol Cell 69, 866–878 e867. 10.1016/j.molcel.2018.02.002.

22. Smeenk, G., and Mailand, N. (2016). Writers, Readers, and Erasers of Histone Ubiquitylation in DNA Double-Strand Break Repair. Front Genet 7, 122. 10.3389/fgene.2016.00122.

23. Soucy, T.A., Smith, P.G., Milhollen, M.A., Berger, A.J., Gavin, J.M., Adhikari, S., Brownell, J.E., Burke, K.E., Cardin, D.P., Critchley, S., et al. (2009). An inhibitor of NEDD8-activating enzyme as a new approach to treat cancer. Nature 458, 732–736. 10.1038/nature07884.

24. Fouad, S., Wells, O.S., Hill, M.A., and D’Angiolella, V. (2019). Cullin Ring Ubiquitin Ligases (CRLs) in Cancer: Responses to Ionizing Radiation (IR) Treatment. Front Physiol 10, 1144. 10.3389/fphys.2019.01144.

25. Vanderdys, V., Allak, A., Guessous, F., Benamar, M., Read, P.W., Jameson, M.J., and Abbas, T. (2018). The Neddylation Inhibitor Pevonedistat (MLN4924) Suppresses and Radiosensitizes Head and Neck Squamous Carcinoma Cells and Tumors. Mol Cancer Ther 17, 368–380. 10.1158/1535-7163.MCT-17-0083.

26. May, D.G., and Roux, K.J. (2019). BioID: A Method to Generate a History of Protein Associations. Methods Mol Biol 2008, 83–95. 10.1007/978-1-4939-9537-0_7.

27. Larochelle, M., Bergeron, D., Arcand, B., and Bachand, F. (2019). Proximity-dependent biotinylation mediated by TurboID to identify protein-protein interaction networks in yeast. J Cell Sci 132. 10.1242/jcs.232249.

28. Fiil, B.K., Damgaard, R.B., Wagner, S.A., Keusekotten, K., Fritsch, M., Bekker-Jensen, S., Mailand, N., Choudhary, C., Komander, D., and Gyrd-Hansen, M. (2013). OTULIN restricts Met1-linked ubiquitination to control innate immune signaling. Mol Cell 50, 818–830. 10.1016/j.molcel.2013.06.004.

29. D’Angiolella, V., Esencay, M., and Pagano, M. (2013). A cyclin without cyclin-dependent kinases: cyclin F controls genome stability through ubiquitin-mediated proteolysis. Trends Cell Biol 23, 135–140. 10.1016/j.tcb.2012.10.011.

30. Walter, D., Hoffmann, S., Komseli, E.S., Rappsilber, J., Gorgoulis, V., and Sorensen, C.S. (2016). SCF(Cyclin F)-dependent degradation of CDC6 suppresses DNA re-replication. Nat Commun 7, 10530. 10.1038/ncomms10530.

31. Klein, D.K., Hoffmann, S., Ahlskog, J.K., O’Hanlon, K., Quaas, M., Larsen, B.D., Rolland, B., Rosner, H.I., Walter, D., Kousholt, A.N., et al. (2015). Cyclin F suppresses B-Myb activity to promote cell cycle checkpoint control. Nat Commun 6, 5800. 10.1038/ncomms6800.

32. Clijsters, L., Hoencamp, C., Calis, J.J.A., Marzio, A., Handgraaf, S.M., Cuitino, M.C., Rosenberg, B.R., Leone, G., and Pagano, M. (2019). Cyclin F Controls Cell-Cycle Transcriptional Outputs by Directing the Degradation of the Three Activator E2Fs. Mol Cell 74, 1264–1277 e1267. 10.1016/j.molcel.2019.04.010.

33. Mavrommati, I., Faedda, R., Galasso, G., Li, J., Burdova, K., Fischer, R., Kessler, B.M., Carrero, Z.I., Guardavaccaro, D., Pagano, M., and D’Angiolella, V. (2018). beta-TrCP- and Casein Kinase II-Mediated Degradation of Cyclin F Controls Timely Mitotic Progression. Cell Rep 24, 3404–3412. 10.1016/j.celrep.2018.08.076.

34. Russo, A.A., Jeffrey, P.D., Patten, A.K., Massague, J., and Pavletich, N.P. (1996). Crystal structure of the p27Kip1 cyclin-dependent-kinase inhibitor bound to the cyclin A-Cdk2 complex. Nature 382, 325–331. 10.1038/382325a0.

35. Okoye, C.N., Rowling, P.J.E., Itzhaki, L.S., and Lindon, C. (2022). Counting Degrons: Lessons From Multivalent Substrates for Targeted Protein Degradation. Front Physiol 13, 913063. 10.3389/fphys.2022.913063.

36. Wheeler, D.A., Takebe, N., Hinoue, T., Hoadley, K.A., Cardenas, M.F., Hamilton, A.M., Laird, P.W., Wang, L., Johnson, A., Dewal, N., et al. (2021). Molecular Features of Cancers Exhibiting Exceptional Responses to Treatment. Cancer Cell 39, 38–53 e37. 10.1016/j.ccell.2020.10.015.

37. Chan, Y.W., and West, S.C. (2018). A new class of ultrafine anaphase bridges generated by homologous recombination. Cell Cycle 17, 2101–2109. 10.1080/15384101.2018.1515555.

38. Chan, Y.W., Fugger, K., and West, S.C. (2018). Unresolved recombination intermediates lead to ultra-fine anaphase bridges, chromosome breaks and aberrations. Nat Cell Biol 20, 92–103. 10.1038/s41556-017-0011-1.

39. Ochs, F., Somyajit, K., Altmeyer, M., Rask, M.B., Lukas, J., and Lukas, C. (2016). 53BP1 fosters fidelity of homology-directed DNA repair. Nat Struct Mol Biol 23, 714–721. 10.1038/nsmb.3251.

40. Zong, D., Chaudhuri, A.R., and Nussenzweig, A. (2016). More end resection is not merrier. Nat Struct Mol Biol 23, 699–701. 10.1038/nsmb.3274.

41. Brambati, A., Sacco, O., Porcella, S., Heyza, J., Kareh, M., Schmidt, J.C., and Sfeir, A. (2023). RHINO restricts MMEJ activity to mitosis. bioRxiv. 10.1101/2023.03.16.532763.

42. Wei, D., Li, H., Yu, J., Sebolt, J.T., Zhao, L., Lawrence, T.S., Smith, P.G., Morgan, M.A., and Sun, Y. (2012). Radiosensitization of human pancreatic cancer cells by MLN4924, an investigational NEDD8-activating enzyme inhibitor. Cancer Res 72, 282–293. 10.1158/0008-5472.CAN-11-2866.

43. Wood, R.D., and Doublie, S. (2022). Genome Protection by DNA Polymerase theta. Annu Rev Genet 56, 207–228. 10.1146/annurev-genet-072920-041046.

44. Harding, S.M., Benci, J.L., Irianto, J., Discher, D.E., Minn, A.J., and Greenberg, R.A. (2017). Mitotic progression following DNA damage enables pattern recognition within micronuclei. Nature 548, 466–470. 10.1038/nature23470.

45. Stricker, S.H., Feber, A., Engstrom, P.G., Caren, H., Kurian, K.M., Takashima, Y., Watts, C., Way, M., Dirks, P., Bertone, P., et al. (2013). Widespread resetting of DNA methylation in glioblastoma-initiating cells suppresses malignant cellular behavior in a lineage-dependent manner. Genes Dev 27, 654–669. 10.1101/gad.212662.112.

46. Chen, E.Y., Tan, C.M., Kou, Y., Duan, Q., Wang, Z., Meirelles, G.V., Clark, N.R., and Ma’ayan, A. (2013). Enrichr: interactive and collaborative HTML5 gene list enrichment analysis tool. BMC Bioinformatics 14, 128. 10.1186/1471-2105-14-128.

47. Huang, D.W., Sherman, B.T., Tan, Q., Collins, J.R., Alvord, W.G., Roayaei, J., Stephens, R., Baseler, M.W., Lane, H.C., and Lempicki, R.A. (2007). The DAVID Gene Functional Classification Tool: a novel biological module-centric algorithm to functionally analyze large gene lists. Genome Biology 8, R183. 10.1186/gb-2007-8-9-r183.

48. Perez-Riverol, Y., Csordas, A., Bai, J., Bernal-Llinares, M., Hewapathirana, S., Kundu, D.J., Inuganti, A., Griss, J., Mayer, G., Eisenacher, M., et al. (2019). The PRIDE database and related tools and resources in 2019: improving support for quantification data. Nucleic Acids Res 47, D442–D450. 10.1093/nar/gky1106.

49. Caron, M.-C., Sharma, A.K., O’Sullivan, J., Myler, L.R., Ferreira, M.T., Rodrigue, A., Coulombe, Y., Ethier, C., Gagné, J.-P., Langelier, M.-F., et al. (2019). Poly(ADP-ribose) polymerase-1 antagonizes DNA resection at double-strand breaks. Nature Communications 10, 2954. 10.1038/s41467-019-10741-9.

